# trFlow: ultrafast all-atom protein conformational ensemble generation via conditional flow matching

**DOI:** 10.64898/2026.10.03.756422

**Authors:** Chenxiao Xiang, Kailong Zhao, Zhenling Peng, Wenkai Wang, Jianyi Yang

## Abstract

Generating diverse, all-atom protein conformations remains challenging, especially for recovering alternative functional states and modeling conformational ensembles. We present trFlow, an ultrafast approach for generating all-atom conformations using geometry-guided flow matching. trFlow leverages three core innovations: (1) a hybrid single- and multi-step sampling scheme that balances accuracy and speed; (2) a decoupled representation-generation architecture that maximizes computational efficiency; and (3) fine-tuning on NMR ensembles to better capture conformational heterogeneity. Extensive benchmarking across dual-conformation proteins, NMR ensembles, and a CASP16 dynamic target shows that trFlow outperforms competing methods in recovering challenging states and reproducing experimental heterogeneity, while being an order of magnitude faster than existing all-atom approaches. These results establish trFlow as a practical and scalable framework for large-scale generation of all-atom conformational ensembles.

## Introduction

Protein structural dynamics are fundamental to biological function [1], as conformational transitions regulate processes such as allostery, ligand recognition, and macromolecular assembly [2]. Accurate structural modeling provides an essential foundation for understanding these processes. Deep-learning methods, exemplified by the AlphaFold series and related approaches, have substantially improved protein structure prediction accuracy [3–5] and expanded modeling to diverse biomolecular complexes [6, 7]. However, these models primarily predict individual high-confidence structures rather than ensembles of interconverting states, limiting their ability to capture conformational heterogeneity [8–10]. Extending structure prediction to functionally relevant conformations is therefore important for understanding protein function.

Several factors make conformations prediction particularly challenging. First, experimental data directly characterizing conformational diversity remain limited. Although the Protein Data Bank (PDB) [11] contains a large number of high-quality structures, only a small fraction of protein sequence clusters contain experimentally resolved, structurally distinct conformations [12], restricting the diversity of states available for model training. Second, the conformations recovered by deep learning-based predictors are strongly influenced by the structural distributions represented in their training data, leading to a bias toward frequently observed states and limiting recovery of less populated or alternative conformations [8]. Consistent with this limitation, the AlphaFold 3 study reported that stochastic sampling across multiple random seeds was insufficient to approximate the conformational ensemble in solution [6]. Finally, molecular dynamics (MD) [13] simulations provide an atomistic framework for exploring protein motions, but their accuracy depends on the underlying force fields, and sufficient sampling of biologically relevant states remains computationally demanding [14]. Efficiently generating ensembles that capture conformational heterogeneity without compromising structural accuracy therefore remains a major challenge.

Several strategies have been explored for multi-conformation prediction. One class of approaches diversifies the inputs to AlphaFold-family models by perturbing multiple sequence alignments (MSAs) [12, 15–17], structural templates [18], or related conditioning information [19, 20]. Although these methods can recover alternative conformations in some cases, their effectiveness depends on informative input variation, and repeated model execution across different perturbations limits large-scale sampling. Recent studies have further shown that MSA clustering does not consistently recover experimentally observed alternative conformations and can even underperform random MSA sampling [21], suggesting that input perturbation alone may be insufficient to overcome the tendency of AlphaFold-like models to favor well-represented structural states [21, 22].

An alternative strategy operates directly on model-predicted geometric representations. Previous studies have shown that predicted inter-residue geometries can encode information associated with alternative conformational states [23]. On this basis, we previously developed trRosettaX2-Dynamics (trX2-D) [24], which heuristically updates predicted two-dimensional geometric restraints to explore alternative conformations without modifying the model input. More recently, SteerAF [25] similarly demonstrated that geometric information encoded in AlphaFold2 distograms can likewise facilitate alternative-state exploration. These studies indicate that predicted geometric distributions provide useful guidance for exploring alternative conformations. However, existing geometry-guided implementations rely on energy minimization or additional gradient-based optimization for structure generation, creating a substantial bottleneck for large-scale sampling.

Recent advances in generative modeling provide a complementary route to protein ensemble generation [26–34]. Diffusion-based models such as BioEmu [29] can efficiently generate large conformational ensembles, but their generative model primarily operates on backbone coordinates, with side-chain conformations reconstructed in a subsequent step. Conditional flow-matching approaches, including AlphaFlow [33] and SimpleFold [30], enable all-atom structure generation, although repeated ensemble sampling remains computationally demanding. AlphaFlow-Lit [31] improves sampling efficiency by decoupling structural representation from generation, albeit with some loss of predictive performance. More broadly, these generative models are not explicitly guided toward specific alternative states; consequently, recovering sparsely populated or structurally distant conformations may require extensive independent sampling. Predicted geometric distributions and generative models offer complementary strengths: the former provide signals for alternative-state exploration, whereas the latter enable flexible and scalable structure generation. Integrating these capabilities may therefore provide a route to more efficient and accurate conformational sampling.

Building on these observations, we present trFlow, an all-atom protein conformations generation framework that integrates geometry-guided conformational exploration with conditional flow matching. Benchmark tests show that trFlow provides an ultrafast and accurate framework for generating all-atom protein conformations.

## Results

### Overview of trFlow

trFlow employs conditional flow-matching (CFM) framework for generating all-atom protein conformations (Fig. 1). As illustrated in Fig. 1a, starting from the input MSA, trRosettaX2 (trX2) [24, 35] predicts inter-residue distance distributions, while AlphaFold2 (AF2) provides a prior structure. A heuristic procedure utilizing iterative single-step CFM sampling rapidly generates an alternative structure from the predicted distance distributions and the prior structure (see Methods and Supplementary Fig. 1a). Both initial structures are used for the subsequent conformation generation step, in which two CFM variants, one pre-trained on X-ray structures and the other fine-tuned on NMR ensembles, generate conformations in parallel. The final predictions are obtained by pooling the results from both variants.

**Fig. 1:**
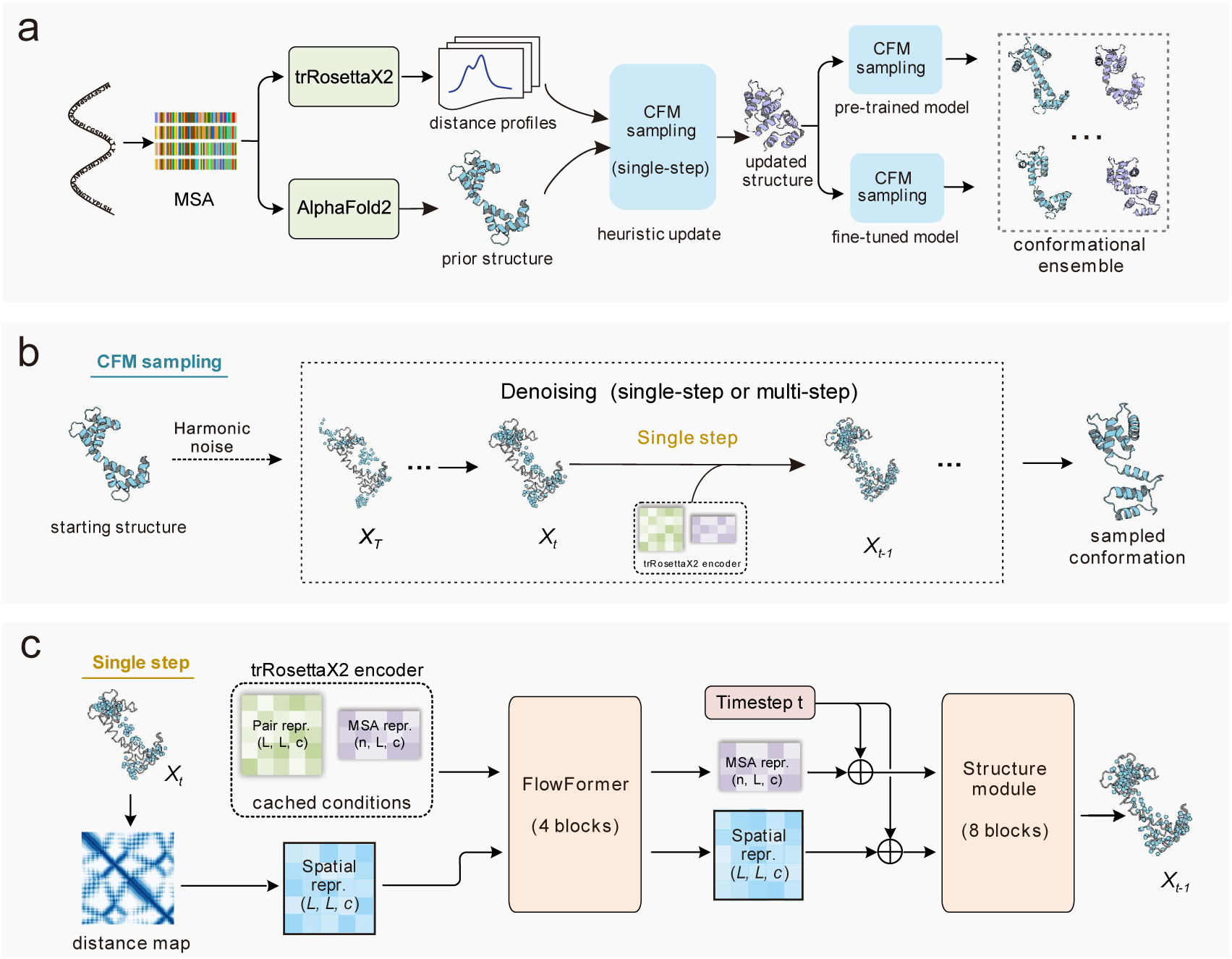
Architecture of trFlow. **(a)** Overall workflow of trFlow. Given an input multiple sequence alignment (MSA), trRosettaX2 predicts inter-residue distance distributions, whereas AlphaFold2 provides the default prior structure. The predicted distance distributions are used in a heuristic geometry-guided conformational exploration procedure based on iterative single-step conditional flow-matching (CFM) generation, yielding an updated structure that serves as an additional sampling prior. The original prior structure and the heuristically updated structure are each used as sampling priors for both the X-ray-pretrained and NMR-fine-tuned model variants, yielding four sampling branches. The ensemble size can be flexibly controlled by varying the number of independent sampling runs performed for each branch. Conformations generated across all branches are pooled to form the final ensemble. **(b)** Schematic of a single CFM sampling run. A noisy structure sampled from the harmonic prior is first aligned to and interpolated with the structural prior to construct the initial flow state, which is then propagated through one or more flow steps to yield a sampled conformation. **(c)** Architecture of the structure generation network used at each flow step. The current flow state is encoded as a spatial representation. This representation, together with the cached conditioning representations and a timestep embedding, is processed by FlowFormer and the structure module to predict an all-atom structure, which is subsequently used to update the flow state.

trFlow adopts a hybrid single- and multi-step CFM sampling scheme (1 or 7 steps; Fig. 1b) to generate conformations efficiently. For each sampling run, either single-step or seven-step sampling is randomly selected with equal probability. To further optimize sampling speed, trFlow implements a decoupled representation-generation architecture (Fig. 1c) that caches and reuses conditioning representations from trX2 across repeated sampling runs. Further details on trFlow’s architecture, training, and inference procedures are provided in the Methods and Supplementary Algorithms 1–3.

### trFlow enables accurate and ultrafast generation of all-atom conformations

We evaluated trFlow on a non-redundant benchmark comprising 32 monomeric proteins, each with two experimentally determined conformations, curated from the Cfold test set [12] and the BioEmu OOD60 benchmark [29] (see Methods). The benchmark features substantial conformational rearrangements, quantified by the RMSD between experimental states (denoted as *D*_conf_), with a mean value of 7.59 Å. Details of these conformation pairs are provided in Supplementary Table 1. On this benchmark, trFlow was compared with three representative methods: AlphaFlow, SimpleFold, and BioEmu. Each method was executed locally using its default settings, producing 200 conformations per target.

RMSD served as the primary metric for structural accuracy due to its sensitivity to both local and domain-level structural displacements. To assess the joint recovery of both experimental states, we defined a dual-state recovery metric, RMSD_dual_, computed by determining, for each experimental state, the minimum RMSD among all generated conformations and then averaging the two minimum values (Eqs. (2) and (3)). On average, trFlow achieved the lowest mean RMSD_dual_ (5.36 Å) on this set (Fig. 2a), outperforming BioEmu (5.45 Å), SimpleFold (5.60 Å), and AlphaFlow (5.98 Å). At the individual-target level, trFlow achieved the lowest RMSD_dual_ among the four methods for 12 of 32 targets, compared with 9 for SimpleFold, 6 for BioEmu, and 5 for AlphaFlow (Supplementary Fig. 2a). Head-to-head comparisons showed that trFlow achieved a lower RMSD_dual_ than each baseline on more than half of the targets (Supplementary Fig. 2b). This advantage stems from geometry-guided exploration of alternative states, with further gains from NMR fine-tuning and hybrid sampling, as supported by the ablation analyses below.

**Fig. 2:**
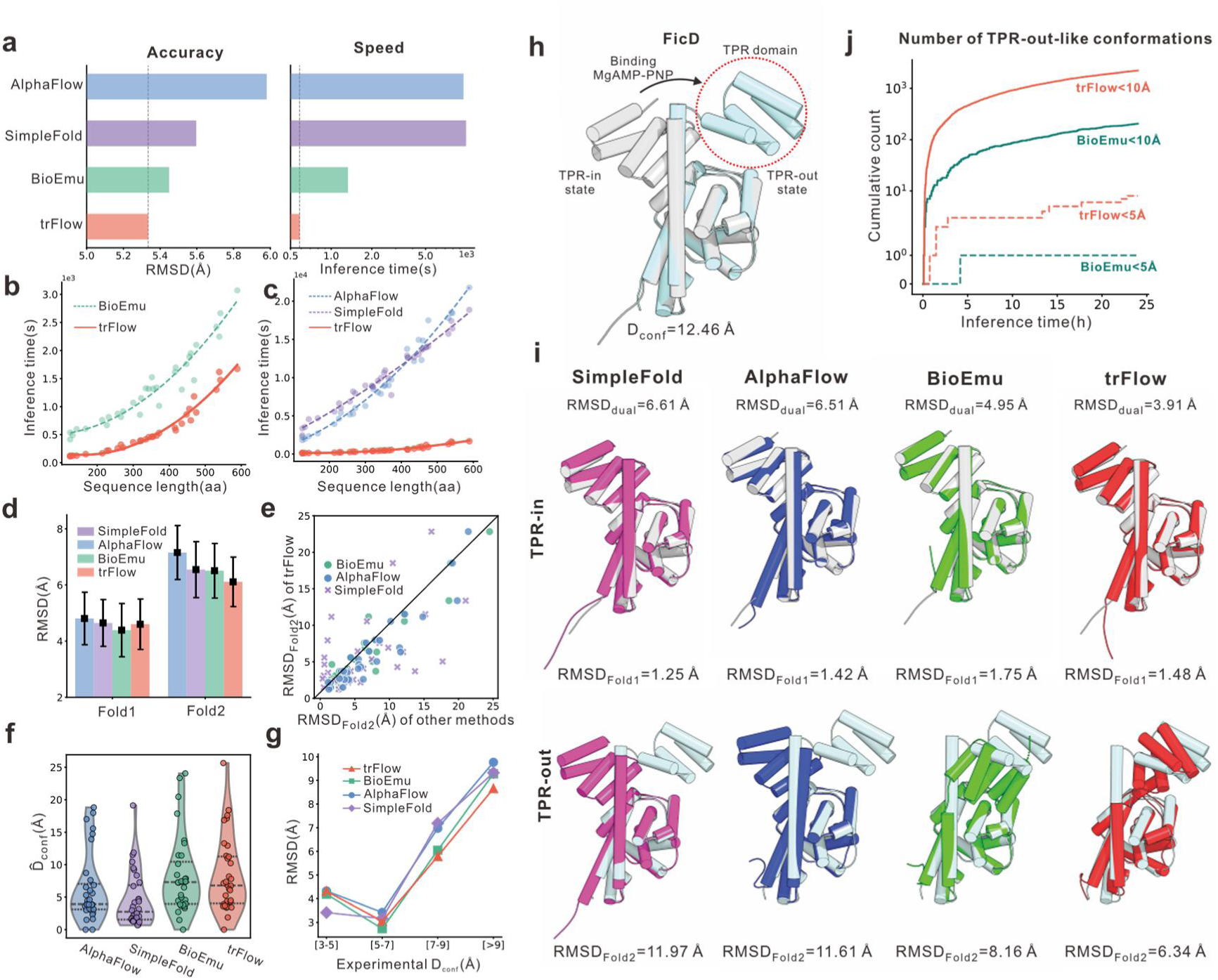
Performance of trFlow on the dual-conformation benchmark. **(a)** Comparison of conformation-recovery accuracy (measured by RMSD_dual_, the average of the minimum RMSDs to the two experimental states) and inference time across the four methods. The reported runtime for trFlow includes the time required to generate the AlphaFold2 prior structure. **(b, c)** Relationship between inference time and sequence length for trFlow and BioEmu (b) and AlphaFlow, SimpleFold (c). Curves represent quadratic fits. **(d)** Recovery of Fold1 and Fold2. Bars show means, and error bars indicate the standard error of the mean across targets. **(e)** Head-to-head comparison of Fold2 recovery between trFlow and the other methods. **(f)** Distributions of 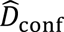 for the four methods; the internal lines represent the quartiles and median. **(g)** Relationship between prediction accuracy and the magnitude of experimentally observed conformational change (*D*_conf_, defined as the RMSD between Fold1 and Fold2). **(h)** Experimental structures of FicD, with TPR-in (Fold1; PDB: 6I7J) shown in gray and TPR-out (Fold2; PDB: 6I7L) in cyan. The TPR domain undergoing the major conformational reorientation is highlighted. **(i)** Representative FicD case study comparing recovery of the large-scale TPR-domain conformational rearrangement by the four methods. Predicted structures are shown in color and the corresponding experimental structures in gray (TPR-in) or cyan (TPR-out). **(j)** Cumulative numbers of FicD conformations generated over 24 hours with RMSD_Fold2_ below 10 Å (solid lines) or 5 Å (dashed lines).

Beyond conformation recovery, trFlow exhibited an even more pronounced advantage in inference efficiency, which is critical for large-scale applications. We benchmarked all four methods on identical hardware using a fixed sampling budget of 200 conformations per target (see Methods for details). Under its default hybrid single- and multi-step sampling configuration, trFlow was the fastest method across all benchmark targets (Fig. 2a, Supplementary Table 2), with an average runtime of 583 seconds per target, i.e., approximately 10 min to generate 200 conformations (including the time required to generate the AF2 prior structure). This corresponds to roughly 2-, 16-, and 17-fold speedups over BioEmu (1,332 seconds), AlphaFlow (9,580 seconds), and SimpleFold (10,010 seconds), respectively. Notably, BioEmu only generates backbone-level conformations, whereas trFlow, AlphaFlow, and SimpleFold generate all-atom structures. Despite this added representational complexity, trFlow remained more than twice as fast as BioEmu and over an order of magnitude faster than other all-atom methods.

To determine if this efficiency advantage persists across larger proteins, we further examined the relationship between inference time and sequence length. Runtime increased strongly with sequence length for all methods (Spearman’s ρ ≥ 0.96), with quadratic fits favored over linear fits (Supplementary Table 3), suggesting superlinear scaling over the tested length range. Despite this shared scaling trend, trFlow maintained a clear speed advantage across all sequence lengths (Fig. 2b-2c). Sequence-length-stratified analysis further confirmed that trFlow was the fastest method in all four length groups (Supplementary Table 4). For proteins longer than 450 residues, trFlow generated 200 all-atom conformations in an average of ∼20 minutes (1,234 seconds), compared with >4 hours for SimpleFold (15,485 seconds) and AlphaFlow (16,899 seconds). This efficiency is achieved by caching the conditioning representations across sampling runs and employing a hybrid structure generation with one or multiple flow steps, thereby reducing repeated representation encoding and network evaluations.

### trFlow improves the recovery of challenging alternative conformations

We next examined the recovery of individual conformational states. For each conformation pair, we classified the two states as Fold1 and Fold2 according to AF2’s prediction preference (see Methods). Fold1 denotes the state more readily predicted by AF2 and thus represents the AF2-preferred state, whereas Fold2 corresponds to the alternative state that is relatively more difficult to recover.

Across all evaluated methods (Fig. 2d), the RMSD to Fold2 (RMSD_Fold2_) was significantly higher than RMSD to Fold1 (RMSD_Fold1_; one-sided Wilcoxon signed-rank test, all P < 0.03), confirming that Fold2 is harder to recover. Although all methods performed similarly on Fold1, trFlow showed a clear advantage for the challenging Fold2 state. Its mean RMSD_Fold2_ was 0.4–1.0 Å lower than those of BioEmu, AlphaFlow, and SimpleFold. Target-wise comparison showed that trFlow achieved a lower RMSD_Fold2_ than each baseline on more than half of the targets, with win rates ranging from 56.3% to 71.9% (Fig. 2e and Supplementary Table 5). These results indicate that the main performance advantage of trFlow lies in the recovery of challenging alternative conformations rather than the relatively accessible AF2-preferred state.

The improved recovery of alternative states was accompanied by greater sampled conformational diversity. We quantified predicted conformational divergence, 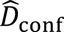 the RMSD between the two generated conformations that best matched Fold1 and Fold2, respectively. As shown in Fig. 2f, trFlow generated more diverse conformations than AlphaFlow and SimpleFold, with a 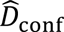 distribution comparable to that BioEmu. It also showed the strongest association with experimental conformational divergence (Spearman’s ρ = 0.62, P < 0.001, compared with 0.41 for BioEmu, 0.28 for AlphaFlow, and 0.16 for SimpleFold; Supplementary Fig. 3).

The greater conformational heterogeneity also translated into improved performance on difficult targets with large experimentally observed conformational heterogeneity (*D*_conf_). For all methods, RMSD_dual_ was positively correlated with *D*_conf_ (Spearman’s *ρ*=0.51∼0.62, all P < 0.005; Fig. 2g and Supplementary Table 6), indicating that larger conformational divergence was generally more difficult to predict. However, trFlow showed the weakest dependence of RMSD_dual_ on conformational divergence, with a linear regression slope of 0.372, compared with >0.48 for other methods (Supplementary Table 6). Consistent with this trend, trFlow achieved the lowest mean RMSD_dual_ in the 7–9 Å bin, and its advantage became more pronounced for targets with *D*_conf_ exceeding 9 Å, confirming its advantage on these challenging targets.

### Case study on FicD illustrates the accuracy and the efficiency of trFlow for large-scale sampling

We examined a representative case, FicD, featuring a large-scale functional conformational rearrangement associated with nucleotide binding in Fig. 2h (PDB: 6I7J/6I7L). The apo Fold1 (6I7J, *TPR*-in) and *MgAMP-PNP-bound* Fold2 (6I7L, *TPR*-out) structures differ by *D*_conf_ = 12.46 Å, primarily involving a near-180° reorientation of the tetratricopeptide repeat (*TPR*) domain [36]. As illustrated in Fig. 2i, all methods readily recovered the more accessible *TPR*-in state, achieving RMSD values of approximately 1–2 Å. In contrast, their performance diverged markedly for the more challenging *TPR*-out state. trFlow achieved the lowest RMSD_Fold2_ of 6.34 Å, compared with 8.16 Å for BioEmu, 11.61 Å for AlphaFlow, and 11.97 Å for SimpleFold, and consequently yielded the lowest RMSD_dual_ of 3.91 Å. Considering that this conformational change is dominated by *TPR*-domain reorientation, which is closely associated with *BiP* recognition and regulation of AMPylation activity [36], we further evaluated recovery of the *TPR*-domain pose after global structural alignment. trFlow again performed best, with a *TPR*-domain RMSD of 8.84 Å, compared with 16.29 Å for BioEmu, 20.17 Å for AlphaFlow, and 20.84 Å for SimpleFold. Therefore, the *TPR*-out-like conformation generated by trFlow more faithfully reproduced both the global topology of the experimental *TPR*-out state and the orientation of its TPR domain.

We further used this challenging target to assess large-scale sampling efficiency. trFlow and BioEmu were run continuously under the same 24-hour computational budget, and we tracked the cumulative yield of *TPR*-out-like conformations generated at different quality thresholds (Fig. 2j). At the RMSD_Fold2_ <10 Å threshold, the two methods showed similar times to first hit, at approximately 6 minutes for trFlow and 5 minutes for BioEmu, indicating that both could rapidly reach the broader *TPR*-out-like conformational region. Their cumulative yields, however, diverged substantially thereafter. Over 24 hours, trFlow generated 2,221 conformations below the 10-Å threshold, compared with 205 by BioEmu. Importantly, trFlow generated only 2.4-fold more conformations during the same period (47,528 versus 20,151), indicating that its >10-fold increase in *TPR*-out-like conformations could not be explained by sampling throughput alone.

The advantage became more pronounced under the stricter RMSD <5 Å criterion: trFlow produced its first qualifying conformation after approximately 45 min, whereas BioEmu required approximately 250 min. By the end of the 24-hour run, trFlow had generated 8 conformations below this threshold, compared with only one for BioEmu. Together, these results show that the sampling advantage of trFlow extends beyond higher throughput, enabling sustained generation of conformations approaching the challenging alternative state.

### Experimentally resolved structural priors enhance alternative-state recovery and conformational exploration

Having established the ability of trFlow to recover challenging alternative conformations from the default AF2 prior, we next examined whether an experimentally determined state could serve as an informative prior for directed sampling toward alternative conformations. For each of the 32 dual-conformation targets, we separately used the experimental Fold1 or Fold2 structure as the sampling prior and evaluated recovery of the opposite state. As shown in Fig. 3a-c, using an experimentally resolved state as the sampling prior improved recovery of the opposite state for most targets, as reflected by lower RMSDs. Compared with the default AF2-prior setting, sampling from Fold2 toward Fold1 yielded a 24.2% reduction in mean RMSD_Fold1_ (from 4.60 to 3.49 Å; Fig. 3a), whereas sampling from Fold1 toward Fold2 reduced the mean RMSD_Fold2_ by 19.0% (from 6.11 to 4.95 Å; Fig. 3b).

**Fig. 3:**
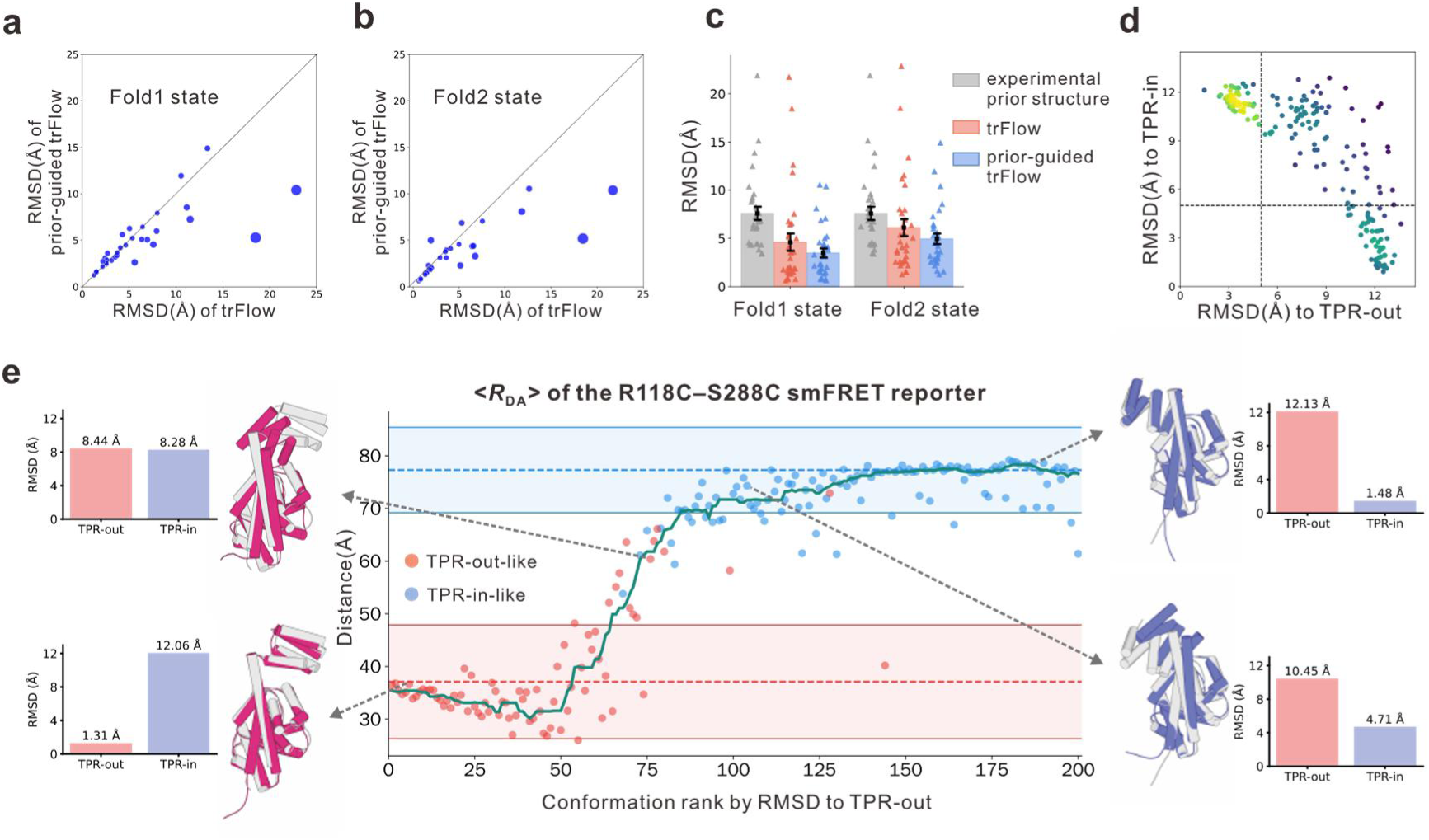
Alternative-state recovery and exploration of intermediate conformational space using experimentally resolved structural priors. **(a, b)** Head-to-head comparison between default and experimental-prior-guided trFlow for recovery of Fold1 (a) and Fold2 (b). Point size is proportional to the perpendicular distance to the diagonal. (**c**) Comparison of state-specific recovery for the input experimental prior, default trFlow (using the AlphaFold2-derived prior), and experimental-prior-guided trFlow. Bars indicate mean values, and error bars denote the standard error of the mean (s.e.m.) across targets. (**d**) Distribution of FicD conformations sampled from the TPR-out structure (PDB: 6I7L) in the two-dimensional space defined by RMSDs to the TPR-out (PDB: 6I7L) and TPR-in (PDB: 6I7J) states. Points are colored according to local sampling density. (**e**) smFRET-based analysis of the FicD conformational series using the R118C-S288C reporter. Conformations are ordered by increasing RMSD from the TPR-out state. The green curve shows the mean donor-acceptor distance (<*R*_DA_>) along the ordered series. Representative conformations and their corresponding RMSDs to the TPR-out and TPR-in states are shown. Red and blue shaded regions indicate the AV-derived reference intervals for the TPR-out and TPR-in states, respectively.

To distinguish genuine exploration toward the target state from simple reproduction of the experimental structure prior, we defined the prior-guided gain (Δ_prior_) as the extent to which the generated conformation moves closer to the target state than the input prior:

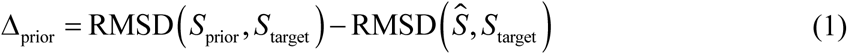

where *S*_prior_ denotes the experimental structure used as the sampling prior, *S*_target_ denotes the opposite experimental state, and *S* is the generated conformation with the lowest RMSD to *S*_target_. A positive Δ_prior_ therefore indicates that trFlow sampled a conformation closer to the target state than the input prior itself, providing evidence of exploration beyond simple prior reproduction. As shown in Fig. 3c and Supplementary Fig. 2c, positive Δ_prior_ values were observed for 29/32 (91%) Fold1-to-Fold2 and 30/32 (94%) Fold2-to-Fold1 tasks, indicating that trFlow generally explored beyond the input experimental structure rather than simply reproducing it.

We then examined whether prior-guided sampling could also support exploration of continuous structural changes between functional states. Using the *TPR*-out structure (PDB: 6I7L) as the sampling prior, the generated conformations extended continuously from the TPR-out-like region toward the *TPR*-in-like region defined by 6I7J in two-dimensional RMSD space (Fig. 3d). We therefore ranked the conformations by their RMSD to 6I7L, producing a series ordered by increasing deviation from the known starting state. The first conformation in this ordering was 0.91 Å from 6I7L, whereas the closest sampled conformation to 6I7J reached 1.48 Å, indicating broad coverage of the structural region spanning the two experimental states.

To determine whether this global structural continuum was accompanied by local rearrangements, we further evaluated five smFRET labeling pairs [37], using the mean donor-acceptor distance (<*R*_DA_>) as an independent local geometric readout (see Methods for calculation details). Among the five labeling pairs, R118C-S288C showed the largest separation between the two experimental endpoints and therefore provided the clearest distinction between the TPR states. Along the RMSD-ordered series, the R118C-S288C donor-acceptor distance shifted overall from the TPR-out-like reference range toward the TPR-in-like reference range despite local fluctuations, with several representative conformations illustrating the accompanying structural progression (Fig. 3e). Given that the donor-acceptor distance was not used for ranking, this coordinated change provides an independent local geometric readout showing that the global TPR reorientation is accompanied by state-specific changes in donor–acceptor distance. The remaining reporters showed weaker or non-monotonic responses along the global RMSD-based ordering (Supplementary Fig. 4). This behavior is consistent with previous smFRET measurements showing reporter- and condition-dependent dynamics beyond those represented by the endpoint crystal structures [37]. These results provide experimental support for the conformational changes captured by trFlow.

### trFlow recapitulates both local flexibility and global diversity in NMR ensembles

We further evaluate trFlow on a benchmark of 77 proteins with experimentally determined NMR ensembles. For each ensemble (predicted or experimental), local flexibility is defined as per-residue root mean square fluctuation (RMSF) across all conformations (Methods, Eq. (4)), while global diversity is defined as the mean pairwise RMSD across all conformations (Methods, Eq. (5)). The local and global accuracy of a generated ensemble is then calculated as the mean absolute error (MAE) relative to the experimental ensemble.

For global accuracy (Fig. 4a), trFlow achieves the lowest median MAE (1.42 Å). This represents reductions of 44.6%, 35.3%, and 83.1% relative to BioEmu, AlphaFlow, and SimpleFold, respectively, indicating that the overall structural spread of the trFlow ensembles most closely matched that of the experimental NMR ensembles. To further examine systematic bias in diversity estimation, we analyzed the signed difference between predicted and experimental ensemble diversity. The bias distribution of trFlow was centered near zero, with a median bias of only +0.02 Å (Supplementary Fig. 5). In contrast, BioEmu showed an overall positive bias, whereas AlphaFlow and SimpleFold showed negative biases. One-sided Wilcoxon signed-rank tests further confirmed that BioEmu significantly overestimated ensemble diversity (P = 0.0073), whereas both AlphaFlow and SimpleFold significantly underestimated it (both P < 0.001). By contrast, the diversity bias of trFlow did not differ significantly from zero (two-sided Wilcoxon signed-rank test, P = 0.829). These results indicate that trFlow provides a more accurate and better-calibrated estimate of experimental ensemble diversity. For local accuracy (Fig. 4b), trFlow achieves the lowest MAE across the benchmark (2.70 Å), compared to 2.82 Å, 3.47 Å, and 6.27 Å for AlphaFlow, BioEmu and SimpleFold, respectively. This indicates that trFlow best captures the residue-level flexibility patterns observed in the experimental ensembles.

**Fig. 4:**
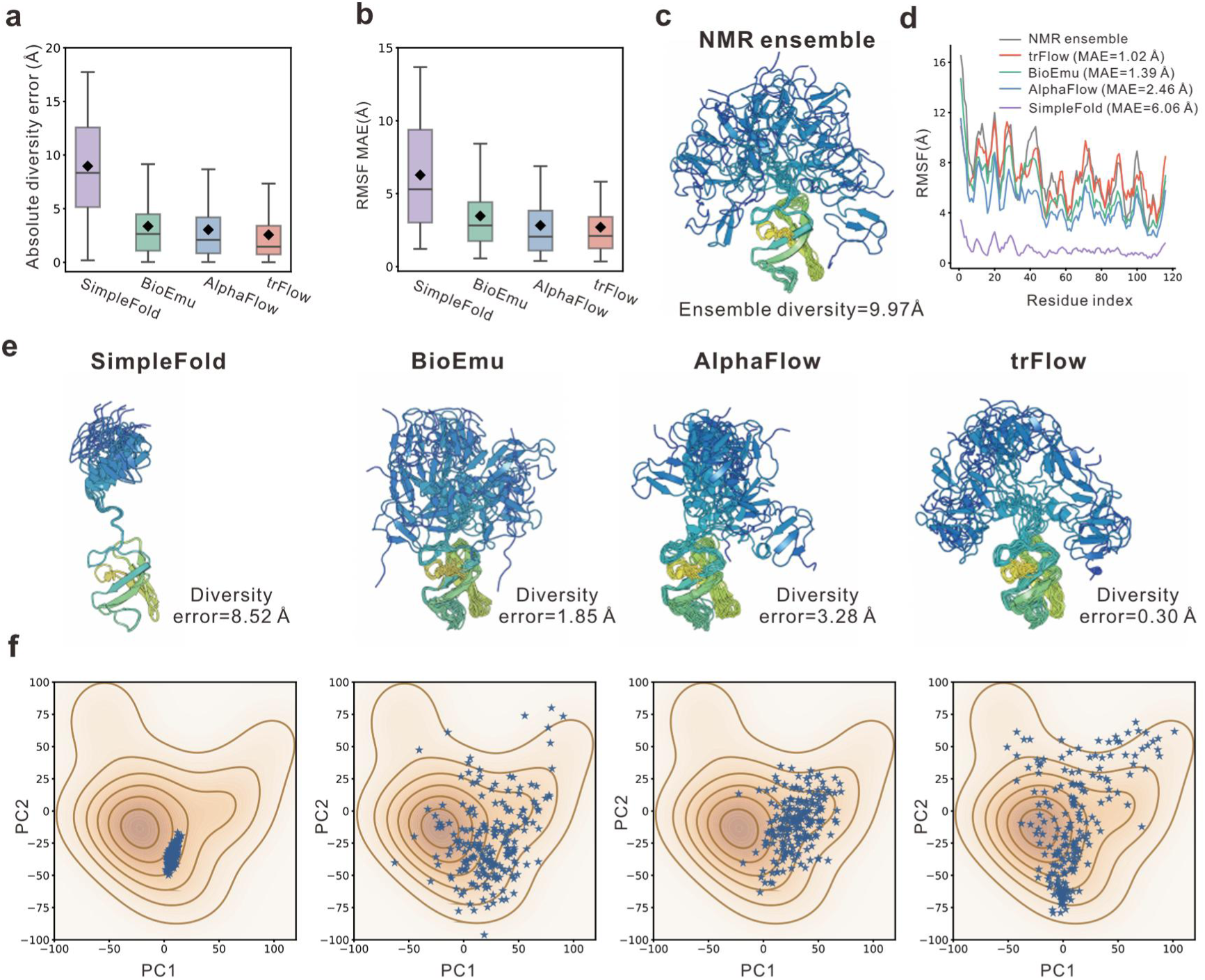
Benchmarking on the NMR ensemble dataset. **(a)** Comparison of global conformational diversity across 77 NMR targets, quantified by the absolute difference between the predicted and experimental mean pairwise RMSDs. **(b)** Comparison of local structural flexibility quantified by MAE between predicted and experimental per-residue RMSF profiles. Black diamonds in (a, b) indicate mean values. **(c)** Experimental NMR ensemble of the representative target PA0128 (PDB: 2AKL). All conformations are superimposed onto the first structure of the NMR ensemble, with the alignment based on the C-terminal SH3-like domain (residues 50-116). Ensemble diversity is defined as the mean pairwise RMSD among conformations. **(d)** Comparison of experimental and predicted per-residue RMSF profiles for 2AKL. The corresponding RMSF MAE values are indicated in the legend. **(e)** Predicted conformational ensembles of 2AKL generated by the four methods. The corresponding diversity errors, defined as the absolute differences between predicted and experimental ensemble diversity, are shown below each ensemble. **(f)** Projection of the predicted ensembles into the principal-component space defined by the experimental NMR ensemble. Orange density contours represent the distribution of experimental NMR conformers, and blue points represent predicted conformations.

To illustrate these performance gains, we examined the NMR ensemble of *Pseudomonas aeruginosa PA0128* (PDB: 2AKL), a protein with two rigid domains connected by a flexible linker that exhibits high conformational diversity in the experimental ensemble (mean pairwise RMSD = 9.97 Å; Fig. 4c). trFlow outperforms all baselines in both local flexibility (RMSF MAE of 1.02 Å vs. 1.39-6.06 Å; Fig. 4d) and global ensemble accuracy (RMSD MAE of 0.30 Å vs. 1.85-8.52 Å; Fig. 4e). Principal component analysis (PCA) further highlights these distinctions (Fig. 4f): SimpleFold and AlphaFlow sampled relatively restricted regions of conformational space, whereas BioEmu sampled more broadly but showed a distribution shifted away from the experimental center. In contrast, trFlow uniquely balanced broad sampling with high fidelity to the central region of the NMR-derived distribution.

### trFlow captures linker-mediated interdomain orientational heterogeneity in CASP16 target T1200

To evaluate whether trFlow can capture interdomain orientation distributions arising from flexible linkers, we applied it to the challenging CASP16 domain–linker–domain (D–L–D) target T1200. This system is derived from the immunoglobulin-binding modules of *Staphylococcus aureus* protein A (SpA) and comprises a relatively rigid ZLBT domain (a B-like domain containing a lanthanide-binding tag) connected by a flexible linker to the wild-type C domain of SpA-N. The linker is thought to constrain the interdomain pose distribution in a manner relevant to antibody binding and SpA-mediated immune evasion [38, 39]. Accordingly, the central challenge is not recovery of a single dominant structure, but reconstruction of the linker-mediated interdomain orientation distribution. Following the CASP16 dynamic-assessment protocol, we back-calculated residual dipolar couplings (RDCs) from the predicted ensembles and compared them with the experimental orthogonal linear combinations (OLCs) of RDCs by linear regression. Pearson correlation, regression slope, and absolute *y*-intercept were used to quantify RDC agreement, while anisotropicity was used to characterize the inferred interdomain orientation distribution. These four metrics were combined into a Chebyshev distance for the final assessment [39].

Among the four methods, trFlow achieved the lowest Chebyshev distance (0.165), compared with 0.254 for AlphaFlow, 0.448 for BioEmu, and 0.999 for SimpleFold (Fig. 5a left). trFlow also outperformed all officially assessed CASP16 T1200 submissions, among which the best value was 0.30 (Fig. 5b). Examination of the individual RDC regression statistics further clarified the basis of this integrated performance (Fig. 5a, right, and Fig. 5c). Although BioEmu achieved the highest Pearson correlation (*r*=0.78), its regression slope of 5.10 deviated substantially from the ideal value of 1.0. This large slope indicates pronounced overestimation of the back-calculated RDC amplitudes and suggests insufficient interdomain orientational heterogeneity, resulting in weak orientational averaging within the predicted ensemble. AlphaFlow and SimpleFold showed similarly large deviations, with slopes of 3.96 and 5.17, respectively. In comparison, trFlow achieved a slope of 1.48, the closest to the ideal value among the four methods. Moreover, trFlow showed a very small absolute intercept (0.22 Hz; compared with 0.78 Hz for BioEmu), indicating minimal systematic offset between the back-calculated and experimental RDC values. Together, these results suggest that the degree of orientational heterogeneity in the trFlow ensemble more closely matches that supported by the experimental RDCs.

**Fig. 5:**
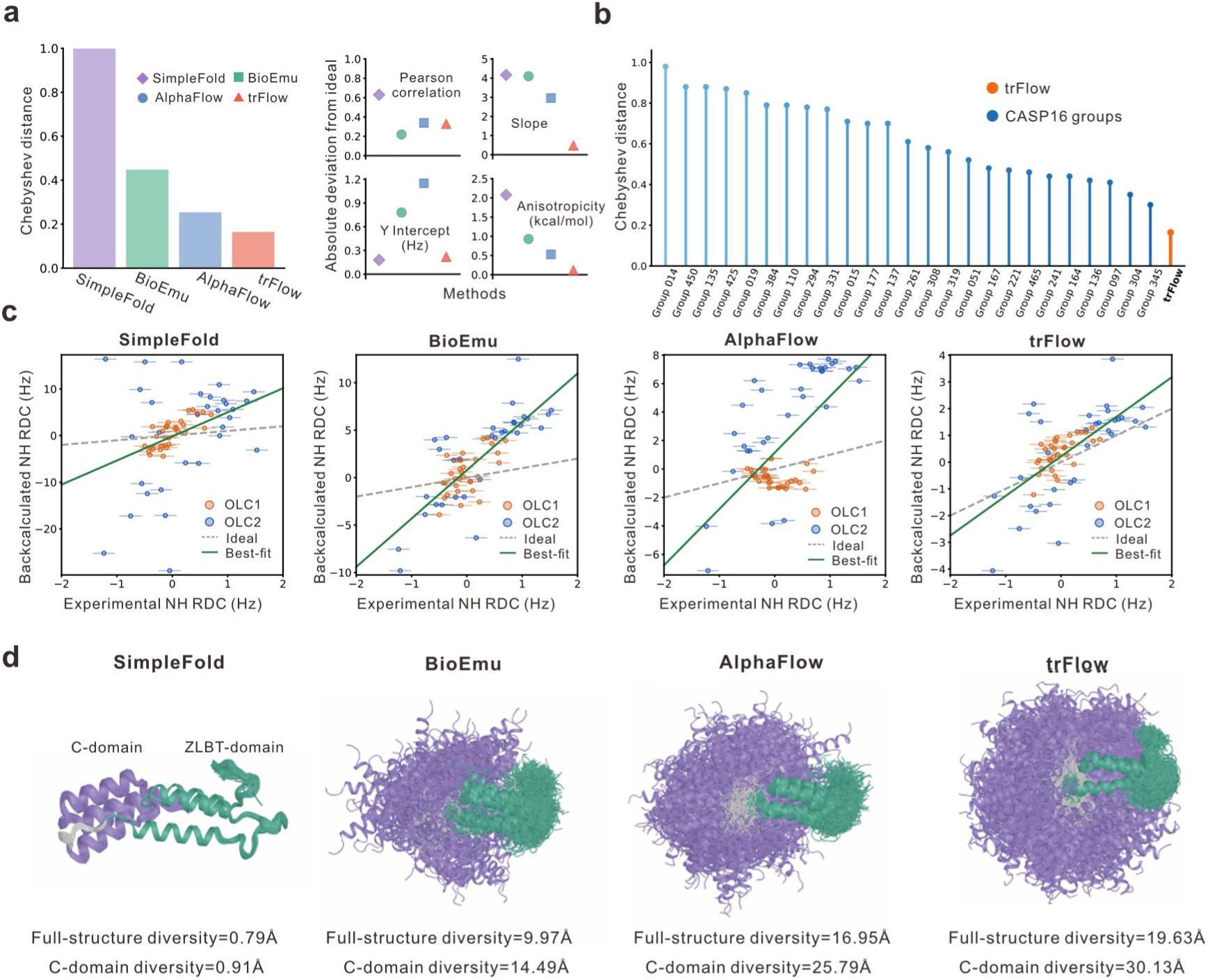
RDC-based evaluation of trFlow on CASP16 T1200. **(a)** CASP16 T1200 assessment of the four methods. Left, overall Chebyshev distance; right, deviations of the four-component metrics from their ideal values or reference ranges. **(b)** Comparison of trFlow with the officially assessed CASP16 T1200 groups ranked by Chebyshev distance. **(c)** Linear regression between ensemble-averaged back-calculated and experimental NH RDCs for OLC1 and OLC2. Dashed gray and solid green lines indicate the ideal relationship and the best-fit regression, respectively. **(d)** Conformational spread of the predicted T1200 ensembles after alignment on the ZLBT domain, illustrating the sampled interdomain positional and orientational variability.

RDC regression statistics quantify agreement with the experimental RDC profile but do not directly characterize the underlying orientation distribution itself. We therefore examined anisotropicity, the fourth component of the CASP16 assessment, which quantifies the deviation of the inferred interdomain orientation distribution from a uniform distribution in free-energy terms. Under the Continuous Distribution of Interdomain Orientations (CDIO) framework [38], the RDC-fitted CDIO solutions span an anisotropicity range of 0.09–0.42 kcal/mol [39]. As illustrated in the right panel of Fig. 5a, trFlow yielded an anisotropicity of 0.54 kcal/mol, the closest to this range among the four methods. In contrast, AlphaFlow, BioEmu, and SimpleFold yielded values of 0.95, 1.35, and 2.50 kcal/mol, respectively. This direct assessment of the orientation distribution further indicates that trFlow more closely reproduced the experimentally supported interdomain orientational heterogeneity of T1200.

To relate these assessment differences to the underlying structural sampling, we next quantified interdomain conformational variability within each predicted ensemble. After aligning all conformations on the ZLBT domain, we calculated the ensemble diversity (mean pairwise RMSD) for both the full structure and the C domain. As illustrated in Fig. 5d, trFlow exhibited the greatest conformational spread, with values of 19.63 Å for the full structure and 30.13 Å for the C domain. The corresponding values were 16.95 and 25.79 Å for AlphaFlow, 9.97 and 14.49 Å for BioEmu, and only 0.79 and 0.91 Å for SimpleFold. Consistent with these quantitative results, trFlow sampled the broadest range of C-domain positions and orientations among the four methods. This broader structural sampling is consistent with the RDC and anisotropicity analyses above and provides a direct structural explanation for the improved agreement of trFlow with the experimental orientational heterogeneity.

In summary, although trFlow was not specifically optimized against RDC data, its generated ensemble showed favorable agreement across multiple dimensions of the CASP16 assessment. Its anisotropicity nevertheless remained slightly above the CDIO-derived reference range, indicating that precise reconstruction of continuous interdomain orientation distributions mediated by flexible linkers remains challenging. Overall, trFlow provides a more experimentally consistent description of linker-mediated interdomain motions, supporting its potential for modeling conformational dynamics in flexible multidomain proteins.

### Ablation study indicates that trFlow benefits from geometry-guided exploration, NMR fine-tuning, and hybrid sampling

To systematically evaluate the contributions of individual components to trFlow performance, we performed ablation analyses on the 32-target dual-conformation benchmark. Three core variants were considered by removing heuristic geometry guidance (*w/o geo*), replacing hybrid single- and multi-step sampling with single-step-only sampling (*w/o multi*), or removing the NMR-fine-tuned model (*w/o NMR*). Each ablation reduced conformation-recovery accuracy (Fig. 6a).

**Fig. 6:**
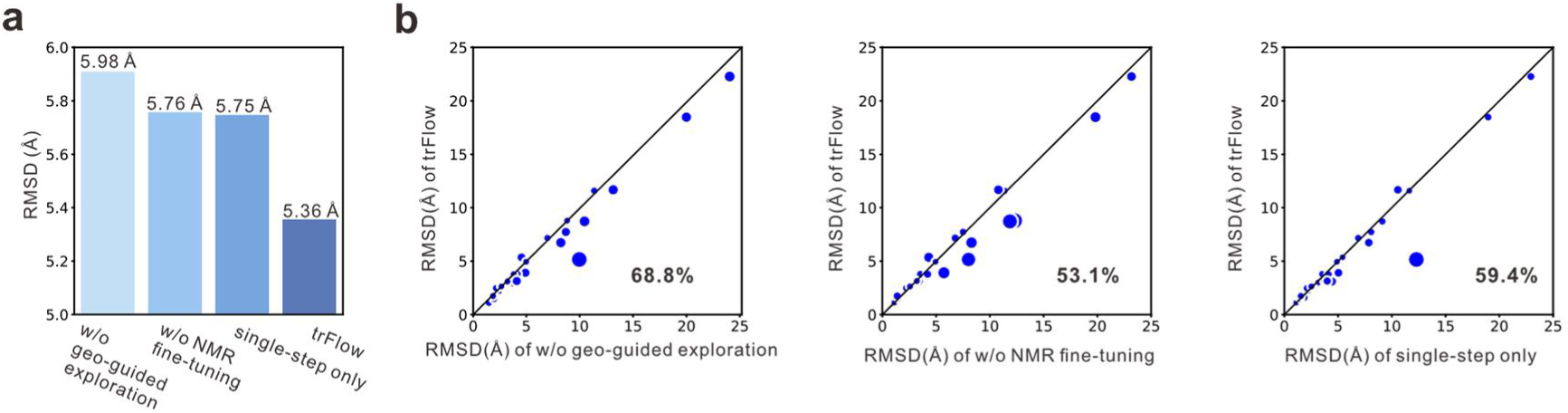
Ablation study of trFlow components. **(a)** Mean RMSD_dual_ of trFlow and the three ablated variants on the dual-conformation benchmark. **(b)** Head-to-head comparisons of RMSD_dual_ between trFlow and each ablated variant. Point size is proportional to the perpendicular distance to the diagonal.

The full model achieved the lowest mean RMSD_dual_ of 5.36 Å. Removing geometry-guided exploration produced the largest degradation, increasing the mean RMSD_dual_ to 5.98 Å (+11.6%), whereas single-step-only sampling and removal of the NMR-fine-tuned model increased it to 5.75 Å (+7.3%) and 5.76 Å (+7.5%), respectively. Head-to-head comparisons in Fig. 6b further confirmed that the full model outperformed each corresponding ablation on most targets. Notably, the contributions of all three components were particularly pronounced for high-difficulty targets (i.e., with *D*_conf_ > 7.59 Å, the benchmark-wide mean). Within this subset, the full model outperformed the corresponding ablated variants on 78.6-85.7% of targets, and reduced mean RMSD_dual_ by 10.5-13.3% relative to the ablations (Supplementary Fig. 6a). By contrast, differences among variants were smaller for low-difficulty targets (Supplementary Fig. 6b), indicating that these components are especially beneficial for proteins undergoing larger conformational rearrangements.

Further analysis revealed distinct roles for the individual components. Removing heuristic geometry-guided exploration produced the largest overall loss in both Fold1 and Fold2 recovery but had little effect on predicted conformational divergence, with the mean 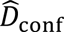 changing only from 8.37 to 8.42 Å (Supplementary Table 7). In contrast, removal of the NMR-fine-tuned model had the smallest effect on RMSD_Fold1_ among the three ablations (from 4.60 Å to 4.69 Å), but produced the clearest reduction in 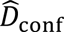 (from 8.37 to 8.05 Å, with decreases observed for 21 of 32 targets). Together with the larger degradation in Fold2 recovery (6.11 Å to 6.83 Å), these results suggest that the NMR-fine-tuned model contributes primarily to maintaining broader separation between sampled states and improving recovery of challenging alternative conformations, rather than the more accessible Fold1 state.

Importantly, the relatively modest accuracy loss associated with single-step-only sampling (a 7.3% increase in mean RMSD_dual_ to 5.75 Å) was accompanied by a 58% reduction in average inference time, to 245 seconds per target (Supplementary Fig. 6c). Multi-step-only sampling, using seven steps per conformation, improved accuracy to 5.60 Å compared to single-step-only sampling but increased the runtime to 844 seconds per target. By comparison, the default hybrid-step strategy achieved a still lower mean RMSD_dual_ of 5.36 Å while requiring 31% less runtime than multi-step-only sampling, thereby outperforming it in both accuracy and efficiency (Supplementary Fig. 6c-d). Overall, the hybrid strategy provided the most favorable balance between conformational recovery accuracy and computational efficiency among the three sampling schemes.

## Discussion

In summary, we have presented trFlow, an ultrafast conditional flow-matching framework for all-atom protein conformational ensemble generation. By integrating geometry-guided exploration, a hybrid single- and multi-step sampling scheme, a decoupled representation-generation architecture, and NMR fine-tuning, trFlow achieves state-of-the-art accuracy in recovering challenging alternative conformations while being up to an order of magnitude faster than existing all-atom methods. Extensive benchmarks across dual-conformation proteins, NMR ensembles, and the CASP16 dynamic target T1200 demonstrate that trFlow not only matches experimental conformational heterogeneity but also provides a practical and scalable solution for large-scale generation. Ablation analyses further confirm the distinct contributions of each component, particularly for proteins undergoing large conformational rearrangements. Looking forward, trFlow offers a promising foundation for integrating experimental restraints and for modeling conformational dynamics in larger and more complex systems, with potential applications in drug discovery, protein engineering, and mechanistic studies of flexible multidomain proteins.

## Methods

### Construction of datasets

To train and evaluate trFlow, we constructed two training datasets and two benchmark datasets. Sequence clustering and redundancy removal were performed using CD-HIT (V4.8.1) [40], ensuring that all benchmark proteins shared <30% sequence identity with proteins in either training dataset.

#### Dual-conformation benchmark

This set was a dual-conformation benchmark comprising 32 proteins, each with two discrete experimental conformations. Candidate pairs were collected from the Cfold test set [12] and the BioEmu OOD60 benchmark [29], yielding 259 conformation pairs in total. To minimize potential training-set overlap and ensure a consistent evaluation setting, we retained only pairs satisfying all of the following criteria: both experimental structures were deposited after the AF2 training cutoff of May 1, 2018; the two conformations shared greater than 90% sequence identity; both conformations corresponded to monomeric proteins; the protein length was below 600 residues; and the observed structural difference was not primarily attributable to crystallization-specific construct engineering, chemical modification, or crystal-packing effects. The resulting benchmark contained 32 conformation pairs, with a mean RMSD of 7.59 Å between the two experimental states.

To label the two experimental states according to AF2 prediction preference, we followed the strategy used in previous studies [22, 24]. For each conformation pair, we calculated the RMSD of each of the five AF2 predictions to both experimental states. The state closer to at least three of the five predictions was designated Fold1, whereas the other was designated Fold2. Fold1 therefore denotes the AF2-preferred state, whereas Fold2 represents the comparatively more challenging alternative state.

#### NMR benchmark set

The second dataset consisted of 77 proteins represented by experimentally determined NMR ensembles, designed to evaluate residue-level flexibility and ensemble-level conformational diversity. Starting from 8,038 monomeric proteins with available NMR ensembles, we first reduced internal sequence redundancy by clustering sequences at 30% sequence identity and retaining one representative per cluster, yielding 3,729 proteins. Proteins sharing more than 30% sequence identity with either training set (see below) were then removed, leaving 2,446 candidates. To focus on proteins exhibiting substantial conformational variation, we retained ensembles with a maximum pairwise RMSD greater than 6 Å and a minimum pairwise TM-score below 0.8. This filtering yielded 881 candidate proteins, of which 20% (176 proteins) were randomly selected for further quality assessment. To reduce the influence of apparent flexibility arising from insufficient restraints, we further assessed whether substantial conformational variation persisted within well-restrained regions, defined as residues participating in at least two NOE restraints with non-neighboring residues (|*i* − *j* ≥ 2). We then recalculated the pairwise RMSDs and TM-scores using only these residues and retained proteins that continued to satisfy the same thresholds. This procedure yielded a final benchmark of 77 proteins exhibiting substantial conformational variation both globally and within experimentally well-restrained regions.

#### X-ray training set

The X-ray training dataset was derived from the 15,051 protein structures originally collected for trRosetta [5]. These structures were non-redundant at 30% sequence identity, released in the PDB before May 1, 2018, resolved by X-ray crystallography, and associated with precomputed multiple sequence alignments (MSAs) containing at least 100 homologous sequences. To avoid overlap with the test sets described above, proteins sharing more than 30% sequence identity with any target in the dual-conformation benchmark were removed. The resulting dataset contained 14,117 structures and was used for model pretraining.

#### NMR training set

The NMR training dataset was constructed from the pool of 8,038 proteins with experimentally determined NMR ensembles described above. Proteins sharing >30% sequence identity with any target in the dual-conformation benchmark were first removed, leaving 7,274 candidate proteins. Because NMR structures are typically deposited as multi-model ensembles, directly treating all deposited conformers as independent training examples would introduce substantial structural redundancy and create large imbalances in the number of conformations contributed by different proteins. We therefore constructed a training set enriched for structural diversity through sequential sequence- and structure-level clustering.

Specifically, the candidate proteins were first clustered at 60% sequence identity, and one representative chain was retained from each cluster, yielding 4,633 proteins. These proteins were then grouped into 3,504 clusters at 30% sequence identity. Within each 30%-identity cluster, all conformers from the corresponding NMR ensembles were pooled and hierarchically clustered according to their pairwise RMSDs. Finally, one representative conformation was retained from each structural cluster. This procedure reduced redundancy among highly similar conformers while preserving structural diversity both within individual NMR ensembles and across homologous proteins. The final dataset contained 12,016 representative structures derived from 4,633 protein chains, with each structure treated as an independent training example during NMR fine-tuning.

### Experimental setup and evaluation metrics

#### MSA generation

For all benchmark proteins, multiple sequence alignments (MSAs) were generated using two independent homologous-sequence search pipelines. The first pipeline used MMseqs2 [41] (v13.45111) to search UniRef30_2202 and the ColabFold environmental sequence database (colabfold_envdb_202108) [42], whereas the second used HHblits (v3.3.0) [43, 44] to search Uniclust30_2018_08 and UniRef30_2023_02. Both pipelines applied an E-value threshold of 0.001, a minimum sequence coverage of 75%, and a maximum sequence identity of 90%. Two MSAs were generated independently for each target, and the MSA with greater sequence depth was selected as the final input for methods that require evolutionary information (including trFlow, AlphaFlow, and BioEmu).

#### Baseline configuration

AlphaFlow, BioEmu, and SimpleFold were selected as generative baselines and evaluated using their officially released implementations with default settings. For the single-sequence model SimpleFold, only the target amino acid sequence was provided as input, and the publicly released SimpleFold-3B model was used. Unless otherwise specified, 200 conformations were generated for each target to construct the evaluation ensemble. All inference experiments were performed on a single NVIDIA A800 GPU. Runtime measurements covered the complete conformation-generation procedure starting from MSA/single sequence.

#### Evaluation metrics

For the dual-conformation benchmark, prediction accuracy was evaluated using RMSD calculated by the TM-score program [45]. Specifically, for each experimentally determined state of a target, the minimum RMSD between that state and all generated conformations was taken as the state-specific recovery error:

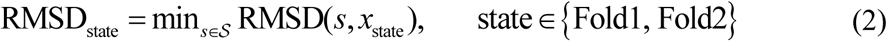

where S denotes the generated conformational ensemble and *x*_state_ denotes the corresponding experimental structure. To jointly evaluate recovery of both experimental states, we further defined RMSD_dual_ as:

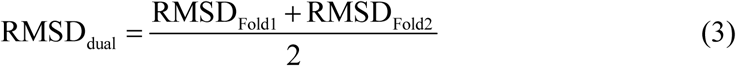

For the NMR ensemble benchmark, we evaluated agreement between predicted and experimental conformational variability at both residue and ensemble levels. Local flexibility was assessed using the mean absolute error (MAE) between the predicted and experimental per-residue RMSF profiles. Ensemble-level diversity was evaluated using the absolute difference between the predicted and experimental mean pairwise RMSDs, termed the absolute diversity error. The corresponding signed difference was used to assess systematic bias, with positive and negative values indicating overestimation and underestimation of ensemble diversity, respectively.

For an ensemble ℰ comprising *M* conformations, the RMSF of residue *i* is defined as:

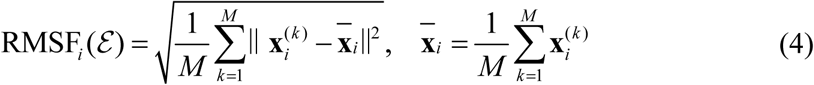

where *x_i_*^(*k*)^ is the aligned coordinate of residue *i* in conformation *k*, and 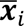 is its mean coordinate across the ensemble. Global ensemble diversity is defined as:

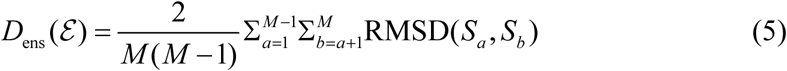

where *S_a_* and *S_b_* denote distinct conformations in the ensemble.

### Network architecture

The core components of the neural network used in trFlow comprise a conditioning representation encoder that extracts MSA-derived features and a structure generation network that generates all-atom conformations. Detailed descriptions of these components are provided below.

#### Conditioning representation encoder

Given an MSA for a target protein, conditioning representations are first extracted using the trRosettaX2 (trX2) encoder [24, 35], which integrates ESM-MSA [46] embeddings, direct coupling analysis (DCA) features, and a stack of trFormer modules. These input features are jointly updated through 12 trFormer blocks [24, 35], producing MSA and pair representations that subsequently serve as conditioning inputs for structure generation. Importantly, the encoder was trained from scratch jointly with the downstream network modules rather than initialized from pretrained trX2 parameters, thereby avoiding potential information leakage from the pretrained trX2 model into the benchmark evaluation.

In addition to the conditioning representations, the trX2 encoder predicts the inter-residue *C_β_* distance distributions. These distributions are subsequently combined with a prior structure, such as an AF2 prediction or an experimentally resolved conformation, during heuristic geometry-guided conformational exploration, yielding an additional structural prior for subsequent ensemble generation (see the section “CFM-based heuristic geometry-guided exploration” for details).

### Structure generation network

To generate structures conditioned on the cached MSA-derived representations, we designed a FlowFormer-based structure generation network (Fig. 1c). For a given intermediate structure, an inter-residue distance matrix is first constructed from its *C_β_* coordinates and embedded into a spatial representation. This spatial representation, together with the cached MSA and pair representations, is then processed by the FlowFormer network.

FlowFormer consists of four stacked FlowFormer blocks (Supplementary Fig. 1b). Each block contains triangle updates, triangle self-attention, a Res2Net module [47], and a feed-forward network. Unlike the AF2 Evoformer, which jointly updates the MSA and pair representations, FlowFormer is designed primarily to update the spatial representation conditioned on the MSA and pair representations. The pair representation remains fixed throughout the FlowFormer blocks, thereby providing persistent inter-residue conditioning information derived from the MSA.

To condition the network on the stage of the flow trajectory, the continuous time variable is encoded separately for the MSA and spatial representations using sinusoidal timestep embeddings and Gaussian Fourier features, respectively. After the FlowFormer updates, the corresponding time embeddings are added to the updated representations. The resulting time-conditioned representations are then passed to the structure module, which adopts the same architecture as that used in trX2 and AF2, to generate the corresponding all-atom three-dimensional structure.

### Model training and ensemble inference

#### Training

For each experimentally determined structure, the corresponding *C_β_* coordinates were extracted and denoted by **x**_exp_. An initial structure **z** was sampled from a harmonic prior and Kabsch-aligned to **x**_exp_. Following EigenFold [26], the harmonic prior was defined as:

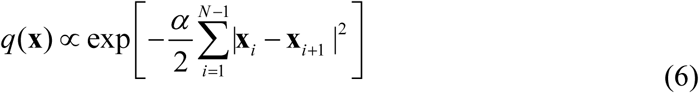

where *N* denotes the protein length and *α*=3/(3.8 Å)^2^ is the harmonic edge strength, chosen to yield a root-mean-square distance of 3.8 Å between adjacent residues. Compared with independent Gaussian noise, this prior introduces correlations between neighboring residues and preserves local chain continuity in the sampled initial structures.

To define a continuous probability path between the experimental and initial structures, intermediate states **x***_t_* were constructed by linear interpolation:

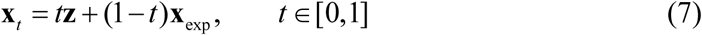

where *t*=0 corresponds to the data endpoint (i.e., the experimental structure) and *t*=1 to the source space (i.e., the initial structure). Rather than predicting the velocity field as in conventional flow matching, trFlow directly predicts the target endpoint structure **x**_exp_ from an intermediate state **x***_t_*. Under the linear interpolation path defined above, endpoint prediction provides an equivalent parameterization of the velocity field, as the corresponding velocity can be analytically recovered from **x***_t_* and the predicted endpoint.

Instead of sampling the continuous time variable *t* uniformly, we adopted a logit-normal sampling scheme inspired by MeanFlow [48]:

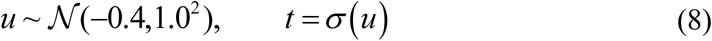

where *σ*(·) denotes the sigmoid function. Compared with uniform time sampling, this distribution biases training toward smaller values of *t*, thereby increasing the frequency of training examples closer to **x**_exp_. This sampling scheme was introduced to strengthen structural recovery under single- and multi-step generation.

To further improve structural recovery under single- and multi-step generation, an auxiliary distance-recovery head was attached to the spatial representation produced by the final FlowFormer block. This head predicts the inter-residue distance distribution and is supervised against the one-hot distance representation derived from the experimental structure using a cross-entropy loss. The overall training objective was defined as

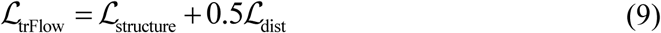

where ℒ_structure_ denotes the structure-prediction loss and ℒ_dist_ denotes the auxiliary distance-recovery loss. ℒ_structure_ follows the original trX2 structure loss (Eq. S1 in SI of [24]), except that the Frame Aligned Point Error (FAPE) term was replaced by its squared form following the AlphaFlow training setting [33].

Training was performed in two stages. trFlow was first trained on the X-ray training dataset described above. Starting from the resulting X-ray-pretrained checkpoint, the entire network was subsequently fine-tuned on the NMR training dataset using the same training objective, yielding the X-ray-pretrained and NMR-fine-tuned models used for ensemble generation. In both stages, the model was optimized using Adam with a learning rate of 0.0001. The complete training procedure is provided in Supplementary Algorithm 1.

#### Ensemble inference

trFlow adopts a decoupled representation–generation architecture to reduce redundant computation during large-scale conformational sampling. For each target, the conditioning representation encoder is executed once to obtain the MSA and pair representations, which are cached and reused across subsequent sampling runs. To introduce stochastic variation into the flow discretization without increasing the number of structure-generation steps, we use stochastic near-uniform time partitioning. For a *K*-step generation process, *K*+1 time intervals are sampled from a symmetric Dirichlet distribution:

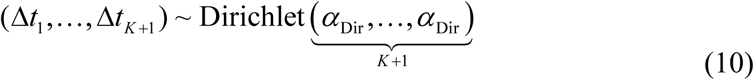

where *α*_Dir_ =5, which concentrates the sampled intervals around uniform spacing while retaining moderate stochastic variation. The corresponding time points are obtained by cumulative summation:

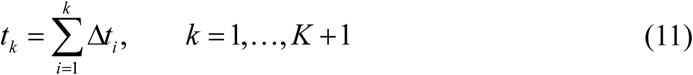

such that *t_K_*_+1_=1.

During inference, each sampling run can be conditioned on a structural prior *S*_prior_ such as an AF2 prediction or an experimentally resolved conformation. The corresponding C*_β_* coordinates are denoted by **x**_prior_. A harmonic-prior structure **z** is first sampled and Kabsch-aligned to **x**_prior_. The initial state of the flow trajectory is then constructed as:

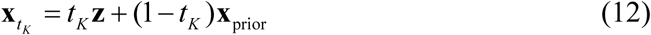

where *t_K_* = 1 − Δ*t_K_* is used as the starting time for generation, rather than the terminal point *t_K_*_+1_=1.

At each time point *t_k_*, the structure generation network takes the current state x*_tk_*, the timestep information, and the cached conditioning representations as inputs to predict an all-atom endpoint structure 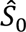. Its C*_β_* coordinates, denoted by 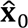 are then used to propagate the current state to *t_k_*_-1_ (Before interpolation, the current flow state x*_tk_* is Kabsch-aligned to the predicted endpoint coordinates, yielding 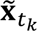.):

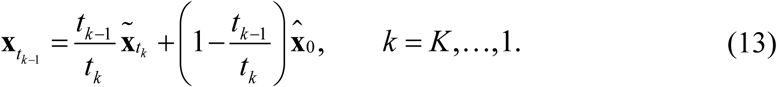

This update follows the same linear interpolation parameterization used during training, progressively propagating the flow state toward the predicted endpoint. After the final step at *t*_0_=0, the all-atom endpoint prediction *Ŝ_0_* is taken as the generated conformation. Unless otherwise specified, the structural prior *S*_prior_ is either the AF2 model_5 prediction or the structure obtained through heuristic geometry-guided exploration.

To balance conformational recovery accuracy and computational efficiency, the default inference protocol uses hybrid single- and multi-step sampling. For each independent sample, either single-step (*K*=1) or seven-step (*K*=7) generation is randomly selected with equal probability. Single-step generation requires only one forward pass through the structure-generation network, whereas multi-step generation iteratively updates the flow state through seven network evaluations.

For final ensemble generation, both the X-ray-pretrained and NMR-fine-tuned models are used. Each model samples from both the AF2 prior and the heuristically updated structural prior, yielding four sampling branches. Conformations generated across all branches are pooled to form the final ensemble. The complete inference procedure is provided in Supplementary Algorithm 2.

### CFM-based heuristic geometry-guided exploration

To exploit alternative-conformation signals encoded in the predicted distance distributions, we adapted the heuristic iterative strategy introduced in trX2-D [24] to the flow-matching framework (Supplementary Fig. 1a). The procedure is initialized with the prior structure and the distance distributions predicted by the trX2 encoder. At each iteration, the current distance distributions are embedded into spatial representations and, together with the cached conditioning representations, provided as conditioning inputs for single-step CFM structure generation. The resulting structure is then used to heuristically update the distance distributions by suppressing probability modes consistent with the current conformation, thereby increasing the relative contribution of alternative distance modes. This process is repeated until the change in the distance distributions between successive rounds falls below a predefined convergence threshold. The final structure is retained as an additional sampling prior and used together with the initial prior during subsequent ensemble generation.

Compared with the original trX2-D implementation, the heuristic geometry-guided exploration procedure in trFlow is redesigned in two key respects. First, heuristic updates are applied only to inter-residue distance distributions, rather than jointly to distance and orientation distributions. Second, the Rosetta-based energy-minimization procedure used for structure generation is replaced by single-step flow-matching generation. These modifications substantially reduce computational cost while preserving the core strategy of geometry-guided exploration toward alternative conformations. The complete procedure is provided in Supplementary Algorithm 3.

### smFRET-based evaluation of FicD conformational series

To evaluate the FicD conformational series generated by trFlow, we analyzed five smFRET labeling pairs reported by Runge *et al.* [37]: L104C–S288C, R118C–S288C, D138C–S288C, K154C–S288C, and S170C–S288C. L104C is located N-terminal to the first α-helix of the TPR domain, R118C within the first TPR motif, D138C in the loop between the two TPR motifs, K154C in the second TPR motif, and S170C near the junction between the TPR domain and the helical linker, whereas S288C is located in the Fic domain. Together, these labeling pairs report distance changes at different positions across the TPR region relative to the Fic domain, providing complementary local measures of TPR-domain reorientation.

For each labeling pair, the mean donor–acceptor distance, 〈*R*_DA_〉, was used as the local geometric observable and was calculated from the accessible-volume (AV) distribution.

The corresponding standard deviation, *σ*_DA_, was used to characterize the spread of the donor–acceptor distance distribution arising from dye accessibility. For each experimental endpoint, the interval 〈*R*_DA_〉 ± *σ*_DA_ was used as the AV-derived reference range for the corresponding TPR state shown in Fig. 3e.

To enable automated AV calculations across the generated ensembles, we used FRETraj v0.2.11 [49] with the three-radii accessible-volume (AV3) model [50]. The C*_β_* atom of each labeled residue was used as the attachment site. We used the same AV geometry parameters as Runge et al.: a linker length of 21 Å, a linker width of 4.5 Å, and AV3 dye radii of 5.5, 4.5, and 1.5 Å for ATTO532 and 7.15, 4.5, and 1.5 Å for ATTO643. The Förster radius for the ATTO532–ATTO643 pair was set to R0=59 Å.

The AV-derived distances reported by Runge et al. were originally calculated using the FRET Positioning and Screening (FPS) software. To verify consistency between the two AV implementations, we recalculated the same experimental endpoint structures used in the published FPS analysis and compared the resulting distance statistics with the reported FPS values. The two implementations showed close agreement in both the mean donor–acceptor distances and the widths of the corresponding distance distributions (Supplementary Table 8), supporting the reliability of the FRETraj-based calculations used here.

### CASP16 T1200 ensemble evaluation

For the CASP16 dynamic target T1200, 300 conformations were generated for each method and used for subsequent ensemble evaluation. For trFlow, structures were generated using the NMR-fine-tuned model with 7 flow steps.

Residual dipolar couplings (RDCs) were back-calculated using the official CASP16 evaluation framework released by the Donald laboratory (https://github.com/donaldlab/CASP16) [39]. Ensemble-averaged back-calculated RDCs were compared with the experimental OLC RDCs by linear regression, yielding the Pearson correlation coefficient (*r*), regression slope, and *y*-intercept. Interdomain orientational anisotropicity was evaluated under the Continuous Distribution of Interdomain Orientations (CDIO) framework [38]. The discrete predicted ensemble was first kernelized into a continuous probability distribution over interdomain orientations using the cdiotools implementation of the CASP16 assessment. Anisotropicity (*A*) was calculated as the free-energy deviation of the predicted orientation distribution *p* from a uniform distribution *p*_unif_ over the three-dimensional rotation space *SO*(3):

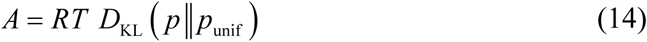

where *D*_KL_ denotes the Kullback–Leibler divergence, *R* is the gas constant and *T*=298.15 K, following the cdiotools implementation. As the integral is evaluated numerically over SO(3), a discrete orientation grid is required. Because the quaternion grid used in the original CASP16 assessment was not distributed with the evaluation resources, we reconstructed a uniform 36,864-point grid using the Hopf-fibration sampling algorithm of Yershova *et al.* [51], which is also adopted in the official assessment framework. The reconstructed grid was then used with the cdiotools [38] implementation to calculate anisotropicity.

The four assessment quantities were scaled according to their ideal and worst possible values defined in the official CASP16 assessment [39]. For the RDC regression statistics, the scaled deviations were

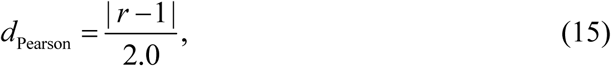

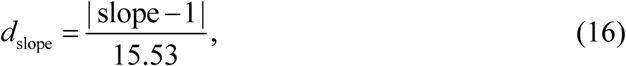

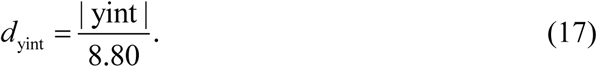

For anisotropicity, the ideal value is defined as the RDC-fitted CDIO interval rather than a single value, and the scaled deviation was calculated as:

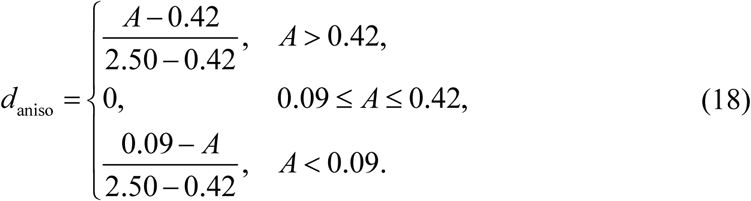

The four scaled deviations were then combined using the four-dimensional Chebyshev distance:

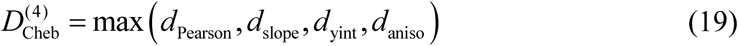

A lower value of 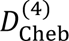 indicates closer agreement with the experimental RDC-based assessment.

## Data availability

All benchmark datasets are available at: https://yanglab.qd.sdu.edu.cn/trFlow/.

## Code availability

The source code is publicly available on GitHub: https://github.com/YangLab-SDU/trFlow.

## Author contributions

J.Y. conceived and supervised the project; C.X. designed and implemented the experiments; W.W. and K.Z. co-supervised the study; Z.P. participated in results analysis. All authors reviewed and approved the final version of the paper.

## Acknowledgements

This work is supported by the National Natural Science Foundation of China (NSFC T2225007, 32430063, 62501364, 62503282), Postdoctoral Fellowship Program and China Postdoctoral Science Foundation (BX20240212 and 2025M783122), the Basic Research Program of Jiangsu Province (BK20250432), and Fundamental Research Funds for the Central Universities.

