## Supplementary Material for "trFlow: ultrafast all-atom protein conformational ensemble generation via conditional flow matching"

### Supplementary Materials

#### Supplementary Algorithms

**Supplementary Algorithm 1** | Training procedure of trFlow.

---

**Algorithm 1** Training

---

**while** not converged **do:**

Sample a training example (MSA, experimental structure $S_{\text{exp}}$)

Extract $C_\beta$ coordinates $\mathbf{x}_{\text{exp}} \leftarrow$ BetaCarbons ($S_{\text{exp}}$)

Sample harmonic-prior noise $\mathbf{z} \sim$ HarmonicPrior ($L$)

Superpose $\mathbf{z} \leftarrow$KabschAlignTo ($\mathbf{z}$, $\mathbf{x}_{\text{exp}}$)

Sample timestep $t \sim \text{LogisticNormal}(-0.4, 1^2)$

Interpolate $\mathbf{x}_t \leftarrow (1 - t) \cdot \mathbf{x}_{\text{exp}} + t \cdot \mathbf{z}$

Construct distance map $\mathbf{D}_t \leftarrow \text{OneHotDistanceBins}(\mathbf{x}_t)$

Predict all-atom structure $\hat{S}_{\text{exp}} \leftarrow$ trFlow (MSA,$\mathbf{D}_t$, $t$)

Update parameters by minimizing $\mathcal{L} = \mathcal{L}_{\text{trFlow}}(\hat{S}_{\text{exp}}, S_{\text{exp}})$

**end while**

---

**Note:** $L$ is the sequence length.

**Supplementary Algorithm 2 |** Structure generation in trFlow.

---

**Algorithm 2** CFM sampling

---

**Input:** MSA, prior structure $S_{\text{prior}}$, Number of steps $K$

**Output:** A sampled all-atom conformation $S_{\text{conf}}$

Encode MSA conditions $\mathcal{C} \leftarrow$ ConditioningRepresentationEncoder(MSA)

Sample $(\Delta t_1, \ldots, \Delta t_{K+1}) \sim$ Dirichlet $(\alpha_{\text{Dir}}, \ldots, \alpha_{\text{Dir}}), \alpha_{\text{Dir}} = 5$

Construct $t_0 = 0, \quad t_k \leftarrow \sum_{i=1}^{k} \Delta t_i, k = 1, \ldots, K+1$

Extract $C_\beta$ coordinates $\mathbf{x}_{\text{prior}} \leftarrow$ BetaCarbons ($S_{\text{prior}}$)

Sample harmonic-prior coordinates $\mathbf{z} \sim$ HarmonicPrior ($L$)

Superpose $\mathbf{z} \leftarrow$ KabschAlignTo ($\mathbf{z}$, $\mathbf{x}_{\text{prior}}$)

$\mathbf{x}_{t_K} \longleftarrow t_K \cdot \mathbf{z} + (1 - t_K)\mathbf{x}_{\text{prior}}$

**for** $k = K, \ldots, 1$ **do**

  Construct distance map $\mathbf{D}_{t_k} \leftarrow$ OneHotDistanceBins($\mathbf{x}_{t_k}$)

  Predict $\hat{S}_0 \leftarrow$ StructureGenerationNetwork ($\mathcal{C}$, $\mathbf{D}_{t_k}$, $t_k$)

  Extract $C_\beta$ coordinates $\hat{\mathbf{x}}_0 \leftarrow$ BetaCarbons ($\hat{S}_0$)

  **if** $k > 1$ **then**

    $\tilde{\mathbf{x}}_{t_k} \leftarrow$ KabschAlignTo ($\mathbf{x}_{t_k}$, $\hat{\mathbf{x}}_0$)

    Interpolate $\mathbf{x}_{t_{k-1}} \leftarrow \frac{t_{k-1}}{t_k} \cdot \tilde{\mathbf{x}}_{t_k} + (1 - \frac{t_{k-1}}{t_k}) \cdot \hat{\mathbf{x}}_0$

  **end if**

**end for**

$S_{\text{conf}} \leftarrow \hat{S}_0$

**Return** $S_{\text{conf}}$

---

**Note:** $L$ is the sequence length.

**Supplementary Algorithm 3 |** CFM-based heuristic geometry-guided exploration.

---

**Algorithm 3** Geometry-guided exploration

---

**Input:** Conditioning representations $\mathcal{C}$, initial distance distributions $\mathbf{D}_0$, prior structure $S_{\text{prior}}$

**Output:** Heuristically updated structure $S_{\text{updated}}$

Initialize $n \leftarrow 0$, $\Delta \leftarrow \infty$, $\epsilon \leftarrow 0.01$, $S_{\text{current}} \leftarrow S_{\text{prior}}$

**while** $\Delta \geq \epsilon$ **do**

  Extract $C_\beta$ coordinates $\mathbf{x}_n \leftarrow \text{BetaCarbons}\ (S_{\text{current}})$

  Construct distance map $\mathbf{D}_n^S \leftarrow \text{OneHotDistanceBins}(\mathbf{x}_n)$

  **for** each residue pair $(i, j)$ **do**

    $\mathbf{p}_n \leftarrow \mathbf{D}_n(i,j)$

    $\mathbf{p}_n^S \leftarrow \mathbf{D}_n^S(i,j)$

    **if** $\|\mathbf{p}_n\|_\infty < 0.5$ **then**

      $\mathbf{p}_{n+1} \leftarrow (f \circ g)(\mathbf{p}_n - 0.5\mathbf{p}_n \odot \mathbf{p}_n^S)$

    **else**

      $\mathbf{p}_{n+1} \leftarrow \mathbf{p}_n$

    **end if**

    Update $\mathbf{D}_{n+1}(\text{i},\text{j}) \leftarrow \mathbf{p}_{n+1}$

  **end for**

  $\Delta \leftarrow \|\mathbf{D}_{n+1} - \mathbf{D}_n\|_\infty$

  $n \leftarrow n + 1$

  **if** $\Delta \geq \epsilon$ **then**

    Generate structure $S_{\text{current}} \leftarrow \text{SingleStepGeneration}(\mathcal{C}, \mathbf{D}_n)$

  **end if**

**end while**

$S_{\text{updated}} \leftarrow S_{\text{current}}$

**Return** $S_{\text{updated}}$

---

**Note:** $\odot$ denotes element-wise multiplication; $g$ denotes normalization; $f$ denotes Gaussian smoothing.

### Supplementary Tables

**Supplementary Table 1 |** Information on the dual-conformation benchmark, including RMSD and TM-score between the Fold1 and Fold2 states.

| Fold1 state | | Fold2 state | | RMSD(Å) between Fold1 and Fold2 | TM-score between Fold1 and Fold2 |
|---|---|---|---|---|---|
| PDB ID | Release Date | PDB ID | Release Date | | |
| 6HKR | 2018-11-28 | 7OXW | 2022-07-13 | 4.37 | 0.76 |
| 6EFR | 2019-01-23 | 7S7V | 2021-10-13 | 4.54 | 0.79 |
| 6NC7 | 2019-04-17 | 6NC9 | 2019-04-17 | 6.82 | 0.65 |
| 6HB0 | 2019-05-01 | 6HBD | 2019-05-01 | 6.03 | 0.72 |
| 6E9N | 2019-05-22 | 6E9O | 2019-05-22 | 4.29 | 0.75 |
| 6HNI | 2019-08-28 | 6HNK | 2019-08-28 | 4.31 | 0.75 |
| 6I7J | 2019-09-25 | 6I7L | 2019-09-25 | 12.46 | 0.79 |
| 6JAQ | 2019-10-30 | 6JAL | 2019-10-30 | 4.43 | 0.75 |
| 6PS5 | 2019-11-13 | 6E67 | 2019-06-05 | 15.11 | 0.64 |
| 6P8R | 2019-12-25 | 6P8O | 2019-12-25 | 6.20 | 0.73 |
| 6O1Z | 2020-03-04 | 6O1X | 2020-02-26 | 7.73 | 0.78 |
| 6L5O | 2020-06-17 | 6L5L | 2020-06-17 | 9.56 | 0.60 |
| 6UHS | 2020-09-16 | 6UHI | 2020-09-16 | 3.81 | 0.67 |
| 6ZJB | 2020-09-16 | 6ZJD | 2020-09-16 | 6.74 | 0.76 |
| 6VGU | 2020-12-16 | 6MFS | 2018-09-26 | 21.88 | 0.50 |
| 6XVI | 2021-01-13 | 6QD6 | 2019-12-18 | 10.34 | 0.75 |
| 6YE8 | 2021-01-20 | 6YED | 2021-01-20 | 4.30 | 0.75 |
| 7LZ9 | 2021-07-14 | 7LZA | 2021-07-14 | 3.37 | 0.72 |
| 7N82 | 2021-07-14 | 6UF2 | 2020-09-30 | 7.61 | 0.46 |
| 7SC3 | 2021-12-01 | 7SC2 | 2021-12-01 | 8.97 | 0.72 |
| 7RDT | 2021-12-29 | 7RDS | 2021-12-29 | 8.75 | 0.54 |
| 7OHG | 2022-06-01 | 6S2U | 2020-07-08 | 7.60 | 0.70 |
| 7XNA | 2022-08-03 | 7XN9 | 2022-08-03 | 8.25 | 0.64 |
| 8DP2 | 2022-08-17 | 7SY9 | 2021-12-15 | 6.00 | 0.74 |
| 8DP6 | 2022-11-09 | 8DP7 | 2022-11-09 | 4.40 | 0.72 |
| 7UY0 | 2022-12-28 | 7UY3 | 2022-12-28 | 6.41 | 0.65 |
| 8BVO | 2023-02-15 | 8BXR | 2023-02-15 | 9.23 | 0.55 |
| 8FT8 | 2023-03-01 | 7N1O | 2021-06-09 | 7.21 | 0.74 |
| 7UGX | 2023-03-29 | 7UH3 | 2023-03-29 | 6.72 | 0.65 |
| 7ZU4 | 2023-05-24 | 7ZU3 | 2023-05-24 | 3.43 | 0.76 |
| 8CRI | 2023-06-07 | 8CRJ | 2023-06-07 | 13.86 | 0.74 |
| 7T4T | 2023-06-14 | 7T4U | 2023-06-14 | 8.06 | 0.69 |

**Supplementary Table 2 |** Inference times of trFlow and the baseline methods on the dual-conformation dataset.

| Sample (Fold1/Fold2) | Sequence length | Inference time (s) for sampling 200 structures per target | | | |
|---|---|---|---|---|---|
| | | **SimpleFold** | **AlphaFlow** | **BioEmu** | **trFlow** |
| 6HKR/7OXW | 143 | 4564.14 | 2331.86 | 598.40 | 125.19 |
| 6EFR/7S7V | 537 | 17418.48 | 18503.53 | 2249.26 | 1619.01 |
| 6NC7/6NC9 | 475 | 14088.28 | 17489.31 | 2100.95 | 898.34 |
| 6HB0/6HBD | 322 | 8772.94 | 8497.22 | 1153.91 | 393.42 |
| 6E9N/6E9O | 460 | 13787.13 | 14853.21 | 1951.54 | 756.64 |
| 6HNI/6HNK | 319 | 8905.65 | 8915.44 | 856.95 | 387.48 |
| 6I7J/6I7L | 343 | 9944.06 | 7891.51 | 1317.29 | 454.60 |
| 6JAQ/6JAL | 416 | 12876.24 | 13761.88 | 1617.02 | 621.51 |
| 6PS5/6E67 | 531 | 15731.70 | 17799.69 | 2423.45 | 1302.71 |
| 6P8R/6P8O | 168 | 4695.60 | 2687.93 | 604.41 | 152.59 |
| 6O1Z/6O1X | 365 | 9919.74 | 8997.80 | 1341.37 | 484.72 |
| 6L5O/6L5L | 372 | 7717.88 | 9757.28 | 1015.73 | 529.12 |
| 6UHS/6UHI | 214 | 5570.18 | 3925.26 | 765.06 | 167.34 |
| 6ZJB/6ZJD | 227 | 6639.26 | 4650.70 | 667.99 | 213.37 |
| 6VGU/6MFS | 435 | 12655.77 | 12160.11 | 1304.61 | 696.89 |
| 6XVI/6QD6 | 542 | 16429.78 | 18439.25 | 2664.92 | 1475.37 |
| 6YE8/6YED | 352 | 10635.50 | 8906.19 | 1525.60 | 462.24 |
| 7LZ9/7LZA | 232 | 6049.40 | 5416.76 | 816.00 | 215.94 |
| 7N82/6UF2 | 125 | 2449.67 | 1809.19 | 414.26 | 122.77 |
| 7SC3/7SC2 | 125 | 3347.25 | 2222.60 | 535.91 | 114.75 |
| 7RDT/7RDS | 257 | 6881.57 | 5185.65 | 875.37 | 256.19 |
| 7OHG/6S2U | 355 | 10954.94 | 9019.20 | 1320.47 | 464.74 |
| 7XNA/7XN9 | 589 | 18859.43 | 21788.92 | 3072.33 | 1674.84 |
| 8DP2/7SY9 | 448 | 13037.72 | 14944.98 | 1760.29 | 862.42 |
| 8DP6/8DP7 | 272 | 7954.51 | 6016.26 | 917.58 | 270.05 |
| 7UY0/7UY3 | 456 | 13632.90 | 12304.38 | 1890.62 | 960.89 |
| 8BVO/8BXR | 300 | 8972.77 | 6304.73 | 1071.78 | 363.16 |
| 8FT8/7N1O | 266 | 8185.46 | 5275.25 | 816.53 | 321.55 |
| 7UGX/7UH3 | 418 | 12600.03 | 12639.25 | 1697.58 | 576.34 |
| 7ZU4/7ZU3 | 132 | 3518.94 | 1959.68 | 491.30 | 124.63 |
| 8CRI/8CRJ | 335 | 9577.04 | 8097.73 | 1328.59 | 416.06 |
| 7T4T/7T4U | 469 | 13932.56 | 14017.51 | 1471.08 | 1183.77 |
| **Average** | **343.8** | **10009.58** | **9580.32** | **1332.44** | **583.40** |

**Supplementary Table 3 | Sequence-length scaling of inference time and comparison of linear and quadratic fits.** Spearman's rank correlation was used to assess the association between sequence length and inference time. Linear and quadratic models were fitted to inference time as a function of sequence length. $\Delta AICc = AICc_{linear} - AICc_{quadratic}$; positive values indicate lower AICc and therefore better support for the quadratic model. Spearman tests were two-sided ($n$=32).

| Method | Spearman's $\rho$ | P value | Linear ($R^2_{adj}$) | Quadratic ($R^2_{adj}$) | ΔAICc |
|---|---|---|---|---|---|
| SimpleFold | 0.974 | $7.16 \times 10^{-21}$ | 0.968 | 0.971 | 1.47 |
| AlphaFlow | 0.986 | $8.35 \times 10^{-25}$ | 0.956 | 0.977 | 19.67 |
| BioEmu | 0.960 | $3.24 \times 10^{-18}$ | 0.884 | 0.935 | 16.72 |
| trFlow | 0.996 | $1.94 \times 10^{-33}$ | 0.843 | 0.964 | 45.30 |

**Supplementary Table 4 | Sequence-length-stratified inference times across methods.** The 32 benchmark targets were stratified into four sequence-length ranges, with eight targets in each group. Values represent the mean inference time required to generate 200 conformations per target. Bold values indicate the shortest inference time within each sequence-length group.

| Sequence-length range (aa) | Mean sequence length (aa) | SimpleFold (s) | AlphaFlow (s) | BioEmu (s) | trFlow (s) |
|---|---|---|---|---|---|
| < 250 | 171 | 4604 | 3125 | 612 | **155** |
| 250-349 | 302 | 8649 | 7023 | 1042 | **358** |
| 350-450 | 395 | 11300 | 11273 | 1448 | **587** |
| > 450 | 507 | 15485 | 16899 | 2228 | **1234** |

**Supplementary Table 5 | Comparison of SimpleFold, AlphaFlow, BioEmu and trFlow on the dual-conformation benchmark.**

| Fold1 state | RMSD against Fold1 state (Å) | | | | Fold2 state | RMSD against Fold2 state (Å) | | | |
|---|---|---|---|---|---|---|---|---|---|
| PDB ID | SimpleFold | AlphaFlow | BioEmu | trFlow | PDB ID | SimpleFold | AlphaFlow | BioEmu | trFlow |
| 6HKR | 0.585 | 0.987 | 2.063 | 1.550 | 7OXW | 4.406 | 3.372 | 2.787 | 3.718 |
| 6EFR | 10.434 | 19.362 | 18.879 | 18.459 | 7S7V | 10.483 | 19.117 | 18.859 | 18.498 |
| 6NC7 | 3.380 | 2.874 | 2.631 | 2.916 | 6NC9 | 3.221 | 4.093 | 3.795 | 2.566 |
| 6HB0 | 5.890 | 3.976 | 3.853 | 3.591 | 6HBD | 0.345 | 2.783 | 1.638 | 2.692 |
| 6E9N | 1.406 | 1.859 | 1.406 | 1.924 | 6E9O | 2.659 | 1.257 | 2.755 | 1.248 |
| 6HNI | 0.289 | 0.653 | 0.687 | 0.617 | 6HNK | 1.138 | 3.900 | 2.624 | 3.499 |
| 6I7J | 1.248 | 1.421 | 1.749 | 1.482 | 6I7L | 11.973 | 11.606 | 8.156 | 6.341 |
| 6JAQ | 0.346 | 0.693 | 0.784 | 0.759 | 6JAL | 4.787 | 3.506 | 3.173 | 2.553 |
| 6PS5 | 15.892 | 10.288 | 12.501 | 12.615 | 6E67 | 6.745 | 10.179 | 12.164 | 10.538 |
| 6P8R | 0.346 | 1.899 | 2.162 | 3.498 | 6P8O | 6.233 | 5.490 | 3.905 | 4.063 |
| 6O1Z | 1.391 | 1.466 | 1.243 | 1.741 | 6O1X | 0.605 | 5.575 | 1.682 | 4.634 |
| 6L5O | 2.880 | 11.852 | 2.870 | 6.411 | 6L5L | 1.621 | 14.820 | 6.993 | 11.156 |
| 6UHS | 3.724 | 1.580 | 1.637 | 1.654 | 6UHI | 0.517 | 1.512 | 1.479 | 1.485 |
| 6ZJB | 0.466 | 1.310 | 1.185 | 1.842 | 6ZJD | 0.528 | 3.288 | 2.197 | 3.112 |
| 6VGU | 1.824 | 3.644 | 3.056 | 4.074 | 6MFS | 21.035 | 19.829 | 18.620 | 13.361 |
| 6XVI | 13.132 | 20.806 | 22.721 | 21.713 | 6QD6 | 16.098 | 21.418 | 24.497 | 22.838 |
| 6YE8 | 0.399 | 0.898 | 0.870 | 0.840 | 6YED | 4.036 | 1.480 | 1.027 | 2.162 |
| 7LZ9 | 1.989 | 4.845 | 4.164 | 5.128 | 7LZA | 0.596 | 4.395 | 4.415 | 5.584 |
| 7N82 | 4.998 | 1.460 | 1.476 | 1.315 | 6UF2 | 5.411 | 7.060 | 7.050 | 6.431 |
| 7SC3 | 12.677 | 8.410 | 10.151 | 5.277 | 7SC2 | 17.704 | 8.172 | 7.517 | 5.038 |
| 7RDT | 1.998 | 1.675 | 1.577 | 1.882 | 7RDS | 8.395 | 7.976 | 7.853 | 8.006 |
| 7OHG | 5.304 | 3.077 | 2.166 | 3.591 | 6S2U | 2.075 | 4.210 | 4.326 | 2.421 |
| 7XNA | 13.597 | 11.763 | 12.252 | 11.824 | 7XN9 | 15.201 | 12.207 | 12.064 | 11.513 |
| 8DP2 | 1.576 | 3.020 | 1.171 | 1.312 | 7SY9 | 5.822 | 1.156 | 2.547 | 2.181 |
| 8DP6 | 0.886 | 0.770 | 0.695 | 0.764 | 8DP7 | 2.996 | 3.270 | 3.086 | 1.499 |
| 7UY0 | 3.416 | 3.167 | 2.577 | 2.020 | 7UY3 | 7.440 | 5.520 | 4.917 | 5.347 |
| 8BVO | 12.461 | 1.125 | 1.674 | 1.925 | 8BXR | 11.146 | 5.834 | 5.960 | 4.275 |
| 8FT8 | 5.416 | 7.939 | 7.372 | 7.516 | 7N1O | 1.063 | 8.575 | 8.563 | 7.933 |
| 7UGX | 2.556 | 3.114 | 3.003 | 4.976 | 7UH3 | 3.102 | 6.222 | 2.671 | 2.604 |
| 7ZU4 | 8.073 | 6.370 | 6.012 | 6.740 | 7ZU3 | 8.560 | 6.632 | 6.368 | 7.582 |
| 8CRI | 0.768 | 0.869 | 0.925 | 0.756 | 8CRJ | 13.695 | 3.057 | 8.022 | 3.710 |
| 7T4T | 9.382 | 10.624 | 5.026 | 6.533 | 7T4U | 9.829 | 11.463 | 6.486 | 6.928 |
| **Average** | **4.648** | **4.806** | **4.392** | **4.601** | **Average** | **6.546** | **7.155** | **6.506** | **6.110** |

**Supplementary Table 6 | Dependence of conformation-recovery error ($\mathbf{RMSD_{dual}}$) on experimental conformational divergence ($D_{\mathrm{conf}}$) on the dual-conformation benchmark.**

| Method | Spearman ρ | P value | Regression slope |
|---|---|---|---|
| SimpleFold | 0.615 | $1.81 \times 10^{-4}$ | 0.601 |
| AlphaFlow | 0.544 | $1.29 \times 10^{-3}$ | 0.489 |
| BioEmu | 0.580 | $5.04 \times 10^{-4}$ | 0.510 |
| trFlow | 0.508 | $2.97 \times 10^{-3}$ | 0.372 |

**Supplementary Table 7 | Effects of trFlow component ablations on conformation recovery and predicted conformational divergence (i.e., $\widehat{D}_{\mathrm{conf}}$) on the dual-conformation benchmark.**

| Model | $\mathrm{RMSD_{dual}}$ (Å) | $\mathrm{RMSD_{Fold1}}$ (Å) | $\mathrm{RMSD_{Fold2}}$ (Å) | $\widehat{D}_{\mathrm{conf}}$ (Å) |
|---|---|---|---|---|
| w/o geometric exploration | 5.98 | 5.09 | 6.87 | 8.42 |
| w/o NMR fine-tuning | 5.76 | 4.69 | 6.83 | 8.05 |
| single-step only | 5.75 | 4.86 | 6.64 | 8.49 |
| trFlow | 5.36 | 4.60 | 6.11 | 8.37 |

**Supplementary Table 8 | Comparison of FRETraj-derived and published FPS donor–acceptor distance statistics for FicD experimental states 6I7J and 6I7K.** Values for FRETraj and FPS are reported as "$\langle R_{\mathrm{DA}}\rangle(\sigma_{\mathrm{DA}})$", in Å. FPS values were taken from Table S3 of Runge *et al.* [1]. Across all five labeling pairs and both experimental states, the absolute differences in $\langle R_{\mathrm{DA}}\rangle$ between FRETraj and FPS were smaller than the corresponding published $\sigma_{\mathrm{DA}}$ values. Differences in $\sigma_{\mathrm{DA}}$ between the two AV implementations were at most 1.2 Å, indicating overall agreement in both mean donor–acceptor distances and the widths of the corresponding distance distributions. Note that the 6I7J and 6I7K structures were used here to match the endpoint structures analyzed in the published FPS calculations.

| Labeling pair | 6I7J | | | | 6I7K | | | |
|---|---|---|---|---|---|---|---|---|
| | FRETraj | FPS | $\lvert\Delta\langle R_{\mathrm{DA}}\rangle\rvert$ | $\lvert\Delta\sigma_{\mathrm{DA}}\rvert$ | FRETraj | FPS | $\lvert\Delta\langle R_{\mathrm{DA}}\rangle\rvert$ | $\lvert\Delta\sigma_{\mathrm{DA}}\rvert$ |
| R118C–S288C | 77.3 (8.1) | 73.9 (7.8) | 3.4 | 0.3 | 37.1 (10.9) | 42.1 (11.2) | 5.0 | 0.3 |
| L104C–S288C | 60.6 (10.0) | 60.1 (10.2) | 0.5 | 0.2 | 58.8 (12.2) | 58.6 (12.4) | 0.2 | 0.2 |
| K154C–S288C | 64.8 (8.4) | 61.5 (7.8) | 3.3 | 0.6 | 50.2 (12.6) | 53.9 (11.4) | 3.7 | 1.2 |
| D138C–S288C | 53.3 (9.9) | 54.3 (9.8) | 1.0 | 0.1 | 61.2 (10.9) | 59.8 (10.9) | 1.4 | 0.0 |
| S170C–S288C | 57.2 (11.1) | 55.3 (10.6) | 1.9 | 0.5 | 63.5 (10.5) | 61.1 (10.5) | 2.4 | 0.0 |

### Supplementary Figures

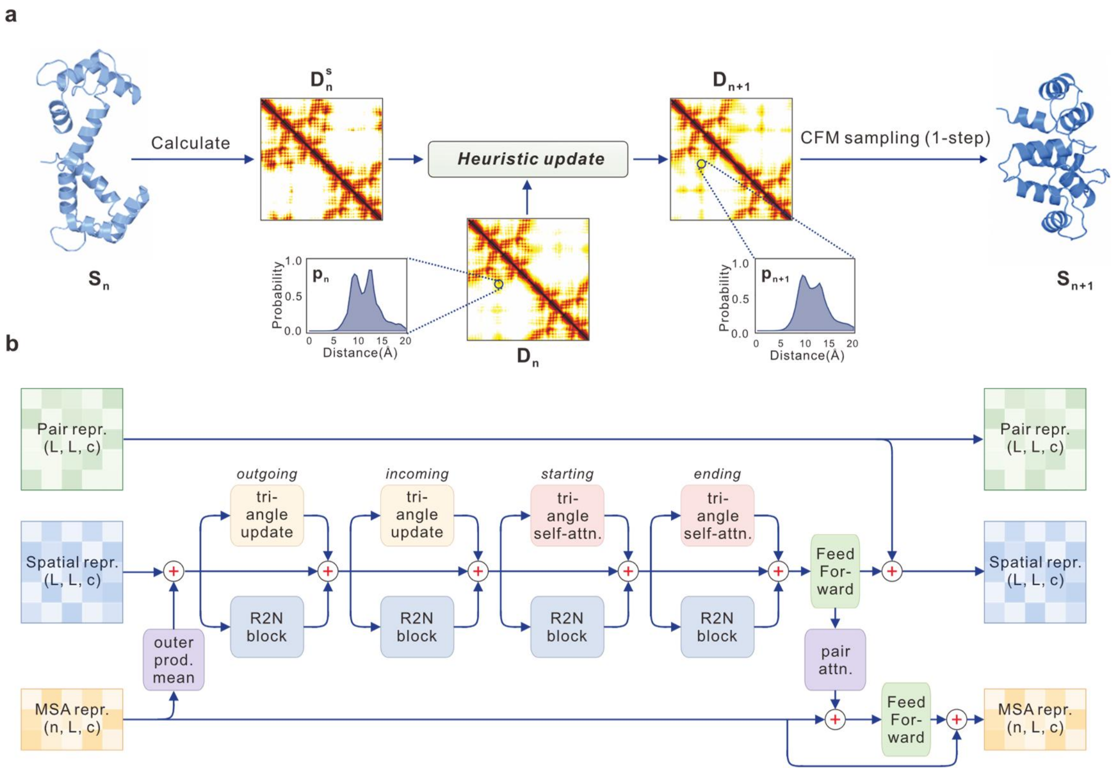


**Supplementary Fig. 1 | Details of geometry-guided conformational exploration and the FlowFormer architecture. (a)** Illustration of one iteration of the heuristic geometry-guided exploration. A structure-derived distance representation, $\mathbf{D}_n^s$, is calculated from the current structure $S_n$ and used to suppress the corresponding probability modes in the current distance distribution $\mathbf{D}_n$, yielding updated distributions $\mathbf{D}_{n+1}$. Single-step CFM sampling conditioned on $\mathbf{D}_{n+1}$ then generates the structure for the next iteration. The procedure is repeated until convergence. **(b)** Architecture of a FlowFormer block. The fixed pair representation serves as conditional information and is repeatedly incorporated into the evolving spatial representation through triangle updates and triangle self-attention, together with Res2Net (R2N) blocks and feed-forward layers. The MSA representation is coupled to the spatial representation through outer-product-mean and pair-attention operations.

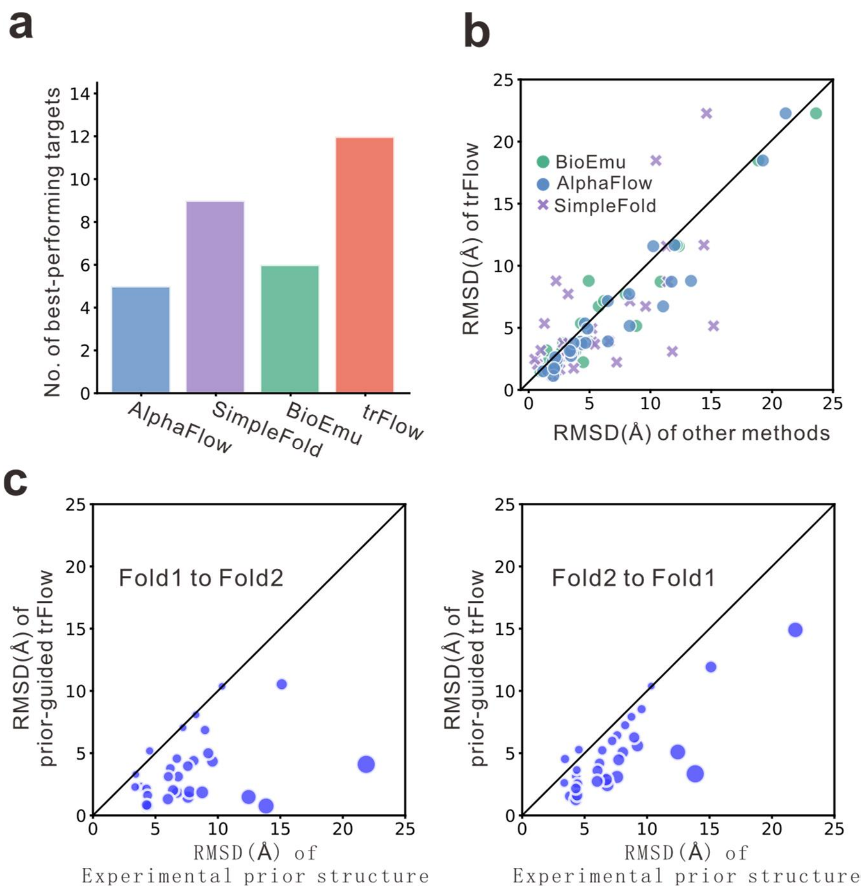


**Supplementary Fig. 2 | Conformation-recovery performance on the dual-conformation benchmark. (a)** Number of targets for which each method achieved the lowest $RMSD_{dual}$. **(b)** Head-to-head comparison of $RMSD_{dual}$ between trFlow and the baseline methods. **(c)** Target-wise comparison between experimental prior-to-target RMSD and the prior-guided trFlow sampling for Fold1-to-Fold2 (left) and Fold2-to-Fold1 (right) tasks. Points below the diagonal indicate positive $\Delta_{prior}$. Point size is proportional to the perpendicular distance to the diagonal.

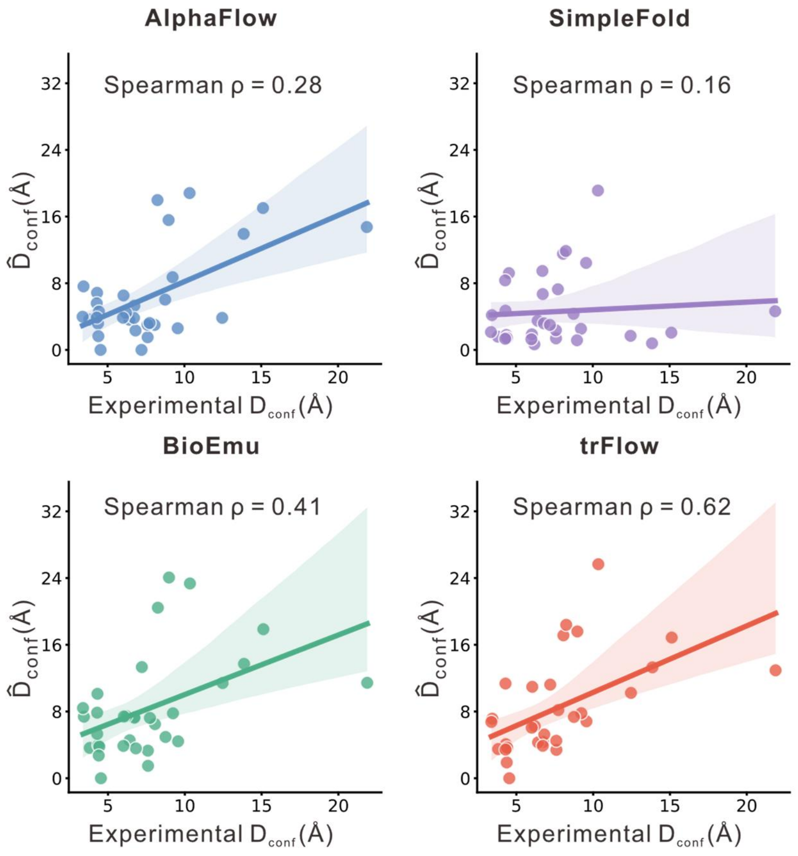


**Supplementary Fig. 3 | Agreement between predicted and experimental conformational divergence.** Each point represents one of the 32 benchmark targets. Lines show linear fits; Spearman's ρ is indicated for each method.

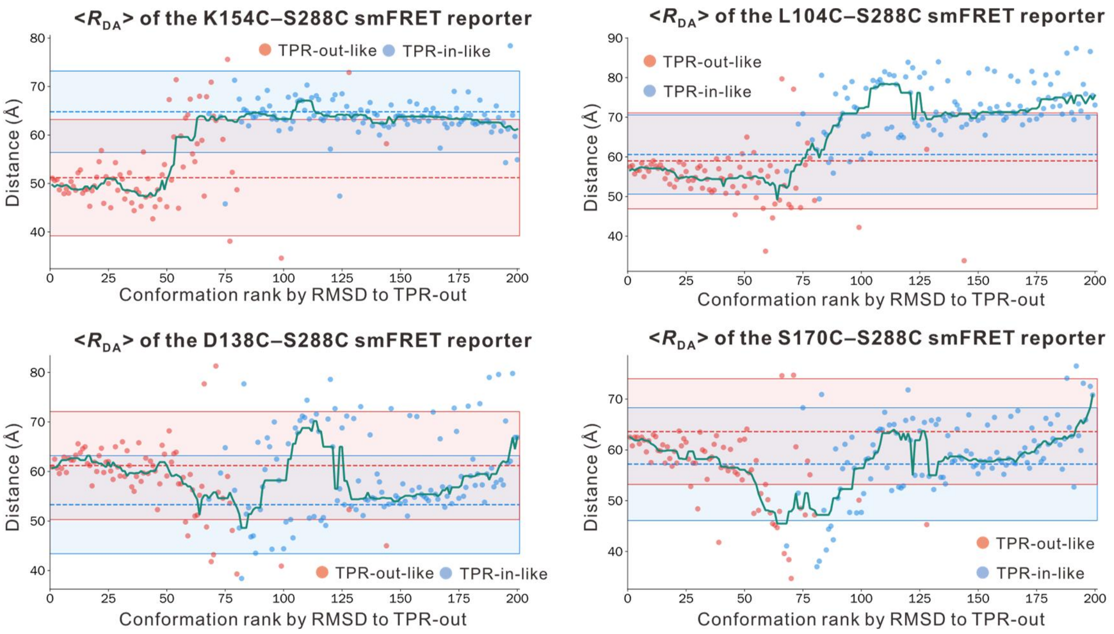


**Supplementary Fig. 4 | smFRET-based analysis of the FicD conformational series using additional labeling pairs.** AV-derived mean donor–acceptor distances (<$R_{DA}$>) for K154C–S288C, L104C–S288C, D138C–S288C, and S170C–S288C along the same conformational series shown in Fig. 3e.

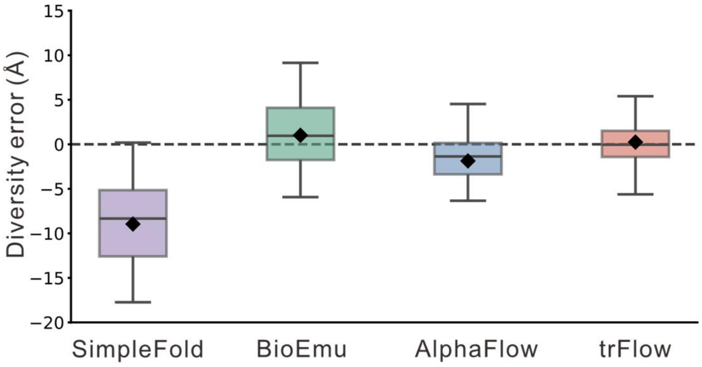


**Supplementary Fig. 5 | Bias in conformational diversity estimation on the NMR benchmark.** Black diamonds indicate mean values, and the dashed line marks zero error.

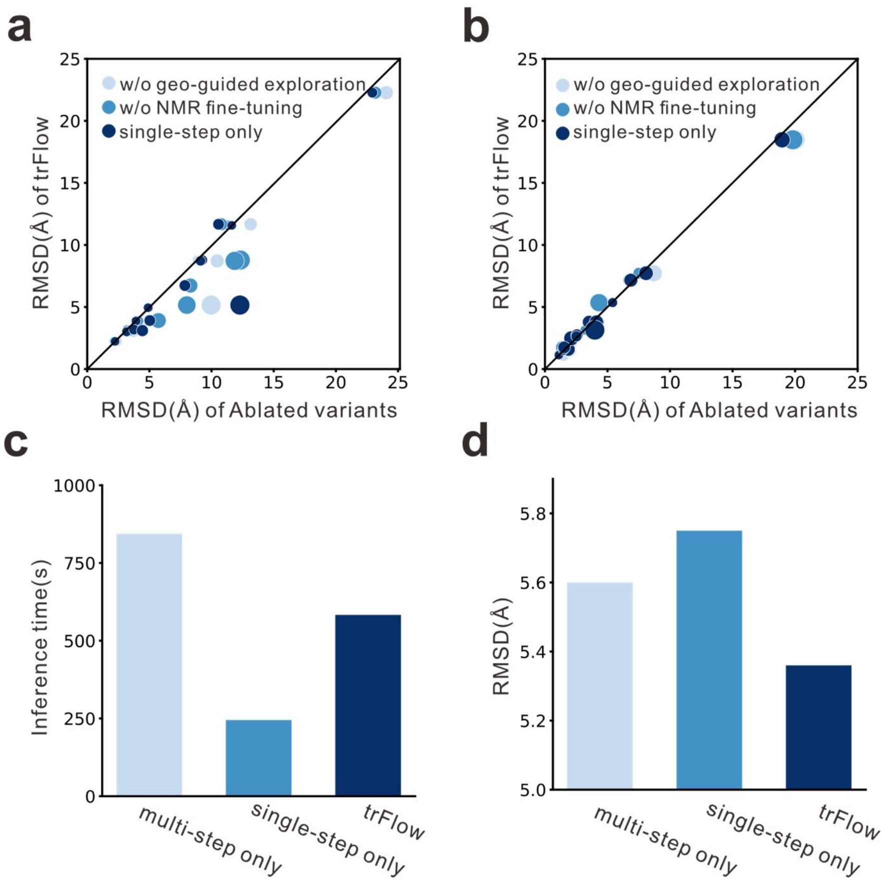


**Supplementary Fig. 6 | Ablation analysis of trFlow components and sampling strategies. (a, b)** Head-to-head comparison of $RMSD_{dual}$ between trFlow and the three ablated variants for high-difficulty ($D_{conf} > 7.59$ Å; **a**) and low-difficulty ($D_{conf} \leq 7.59$ Å; **b**) targets. Points below the diagonal favor trFlow. Point size is proportional to the perpendicular distance to the diagonal. **(c, d)** Comparison of inference time **(c)** and $RMSD_{dual}$ **(d)** among multi-step-only (7 steps), single-step-only, and the default hybrid single/multi-step sampling strategies.
